# The Taste of Blood in Mosquitoes

**DOI:** 10.1101/2020.02.27.954206

**Authors:** Veronica Jové, Zhongyan Gong, Felix J.H. Hol, Zhilei Zhao, Trevor R. Sorrells, Thomas S. Carroll, Manu Prakash, Carolyn S. McBride, Leslie B. Vosshall

## Abstract

Blood-feeding mosquitoes survive by feeding on nectar for metabolic energy, but to develop eggs, females require a blood meal. *Aedes aegypti* females must accurately discriminate between blood and nectar because detection of each meal promotes one of two mutually exclusive feeding programs characterized by distinct sensory appendages, meal sizes, digestive tract targets, and metabolic fates. We investigated the role of the syringe-like blood-feeding appendage, the stylet, and discovered that sexually dimorphic stylet neurons are the first to taste blood. Using pan-neuronal GCaMP calcium imaging, we found that blood is detected by four functionally distinct classes of stylet neurons, each tuned to specific blood components associated with diverse taste qualities. Furthermore, the stylet is specialized to detect blood over nectar. Stylet neurons are insensitive to nectar-specific sugars and responses to glucose, the sugar found in both blood and nectar, depend on the presence of additional blood components. The distinction between blood and nectar is therefore encoded in specialized neurons at the very first level of sensory detection in mosquitoes. This innate ability to recognize blood is the basis of vector-borne disease transmission to millions of people world-wide.

## INTRODUCTION

Animals actively obtain energy from nutrients in food like protein and carbohydrates, which are distinguished by their savory (“umami”) or sweet taste, respectively (Liman et al., 2014; Yarmolinsky et al., 2009). These two taste qualities signal different nutritional values, and animals use diverse strategies to prioritize ingestion of the food source that best matches their current metabolic requirements. For feeding specialists, discrimination between savory and sweet tastes can be hardwired into the animal’s genetic code. Cats are obligate carnivores that have lost the canonical sweet taste receptor but retain a functional umami receptor (Li et al., 2005). Hummingbirds, which are nectar-feeding specialists, have evolved a novel sweet taste receptor from the ancestral umami receptor (Baldwin et al., 2014). For feeding generalists like flies, rodents, and humans, both protein and carbohydrates are useful energy sources and these animals can detect both savory and sweet tastes. Detection of either taste typically promotes feeding unless an animal becomes deficient in a specific nutrient (Deutsch et al., 1989; Leitao-Goncalves et al., 2017; Liu et al., 2017; Murphy et al., 2018; Ribeiro and Dickson, 2010; Simpson et al., 2015; Steck et al., 2018; Vargas et al., 2010). After days of protein deprivation, for example, animals can still detect savory and sweet, but savory taste circuit sensitivity is increased to promote a protein-specific appetite (Liu et al., 2017; Steck et al., 2018). Intrinsic indifference to a taste is ideally suited for specialists that utilize only one food source while acute neuromodulation of taste preference is an effective means for generalists to conditionally prioritize one food source. However, female blood-feeding mosquitoes are specialists with two parallel specific appetites for protein and carbohydrates that each require a different feeding program and fulfil distinct physiological processes. The mechanism that enables mosquitoes to engage mutually exclusive feeding programs for each nutrient is unknown.

The specialized feeding demands of blood-feeding mosquitoes, including *Ae. aegypti*, are linked to nutritional value. Although carbohydrates supplied from nectar are sufficient for energy metabolism in both females and males, protein obtained from blood is necessary for females to develop eggs and successfully reproduce. Mosquitoes take nectar from plant sources like flowers and are likely attracted by olfactory floral cues (Lahondere et al., 2020; Van Handel, 1972). A blood meal must be obtained from a human or other vertebrate animal and *Ae. aegypti* integrate sensory cues like carbon dioxide (CO_2_), heat, and odor to locate their victim (Dekker et al., 2005; Liu and Vosshall, 2019; McMeniman et al., 2014; Takken and Kline, 1989). To procure necessary nutrients from these distinct food sources, females employ two behaviorally and anatomically distinct feeding programs: blood-feeding and nectar-feeding. Each feeding program is linked to a distinct feeding appendage, meal size, and digestive tract (Gordon and Lumsden, 1939; Trembley, 1952). Nectar is detected by the labium (Sanford et al., 2013). Blood is likely detected by the stylet, which pierces skin and directly contacts blood (Gordon and Lumsden, 1939; Trembley, 1952). We use the term “stylet” to refer to the needle-like feeding tube, also known as the labrum, which is used to ingest blood (Lee, 1974). Females typically take small nectar meals but engorge on blood, consuming a volume that reliably doubles their body weight and provides sufficient protein to allow them to produce 100 – 150 eggs per blood meal. Finally, the nectar meal is routed initially to the crop, whereas ingested blood entirely by-passes the crop and is directed to the midgut, which is specialized to digest protein for egg production (Gordon and Lumsden, 1939; Trembley, 1952). Thus, the mosquito has parallel feeding pathways for nectar and blood from the sensory periphery, to visceral organs, to the ultimate metabolic function of the meal. This strict separation in feeding programs may allow the female to maintain a hunger for blood even after taking a nectar meal to sustain her metabolism. From a global health perspective, understanding how the female distinguishes blood from nectar is critical because blood-feeding behavior is required for vector-borne disease transmission and species propagation.

The ability of a female to distinguish between blood and nectar is finely tuned but the underlying mechanism remains unclear. In the absence of human sensory cues like heat and CO_2_, female mosquitoes readily ingest nectar via the nectar-feeding program. In the presence of human sensory cues, females will reliably bite and feed on warm blood delivered in an artificial feeder (Bishop and Gilchrist, 1946; McMeniman et al., 2014). But if the blood meal is replaced with nectar sugars, females reject the meal entirely even though heat and CO_2_ are present (Bishop and Gilchrist, 1946). Therefore, the mechanism that distinguishes between blood and nectar must be flexible enough to promote ingestion of nectar only when a mosquito intends to feed on nectar and not when she intends to feed on blood. A hint may lie in the fact that different sensory neurons are in contact with the meal during the blood- and nectar-feeding programs. If the preference for blood is hardwired into the sensory appendage involved in blood-feeding, we would expect it be a specialized blood detector that is either intrinsically insensitive to nectar sugars, or able to detect nectar sugars differently than the sensory neurons involved in nectar-feeding. Alternatively, blood-feeding and nectar-feeding neurons do not have to be specialized and could have the capacity to detect both blood and nectar. If so, the presence of human cues could increase the sensitivity for blood and/or decrease the sensitivity for nectar sugar to selectively promote blood-feeding. To distinguish between these possibilities, a fundamental understanding of blood and nectar detection in *Ae. aegypti* is crucial.

The sensory mechanisms of blood recognition prior to initiating blood-feeding behavior are unknown. Classic behavioral experiments have demonstrated that the nutritional value of blood as a protein source can be decoupled from blood-feeding behavior. The protein fraction of blood is neither sufficient nor necessary to trigger feeding, but a mixture of key plasma components such as adenosine triphosphate (ATP), sodium chloride (NaCl), and sodium bicarbonate (NaHCO_3_) reliably induces blood-feeding behavior (Galun et al., 1963; Galun et al., 1984; Hosoi, 1959). Importantly, non-hydrolyzable analogues of ATP in saline are still sufficient to trigger engorgement, indicating that energy release from ATP hydrolysis is not required (Galun et al., 1985). Together these results suggest that chemosensory detection of specific blood components is critical for blood recognition.

The stylet is the only sensory appendage that directly contacts blood and is therefore likely the primary structure that evaluates blood prior to initiation of blood-feeding. Electron microscopy studies have revealed the presence of female-specific sensory sensilla at the tip of the stylet (Lee, 1974). Sensilla are specialized insect cuticle structures that house sensory neuron dendrites. Chemical ligands enter chemosensory sensilla through pores to directly contact these dendrites (Stocker, 1994). Extracellular recordings from one stylet sensillum type documented neuronal activity in response to specific plasma components (Werner-Reiss et al., 1999a,b,c). In the two decades since these studies were reported, many questions remain. Do individual stylet sensory neurons respond to whole blood as a mixture or are they are tuned to recognize individual blood components? Blood contains components that are traditionally associated with distinct taste qualities including sodium chloride (salty), protein (umami), glucose (sweet), and CO_2_ (sour/carbonation). Is blood recognized as a single taste quality, or are multiple taste qualities integrated to form the perception of blood? This work set out to discover how mosquitoes perceive the taste of blood and how they distinguish it from the taste of nectar.

Here we show that female *Ae. aegypti* mosquitoes possess sexually dimorphic stylet neurons that are specialized to distinguish blood from nectar. Using pan-neuronal GCaMP calcium imaging, we found that stylet neurons robustly respond to blood and its components, but are insensitive to nectar-specific sugars. We defined a mixture of four blood components—ATP, glucose, sodium bicarbonate, and sodium chloride—that reliably trigger blood-feeding behavior and determined that these ligands activate the same population of stylet neurons as blood. By presenting these ligands individually or as mixtures, we show that the taste of blood is combinatorial across multiple taste qualities. We defined functionally distinct subsets of stylet sensory neurons that are selectively tuned to specific blood components. Since the transcriptional profile of stylet neurons was unknown, we performed RNA-seq on the stylet to identify genetic markers that selectively label these neuronal subsets. We identified *Ir7a* and *Ir7f* as female stylet-specific transcripts and generated driver lines for both genes using CRISPR-Cas9 genome editing. We found that each driver line labels a functionally distinct subset of blood-sensitive stylet neurons activated by different components of blood. Finally, we discovered polymodal stylet neurons that respond to physiological levels of blood glucose only in the presence of additional blood components: sodium chloride and sodium bicarbonate. Importantly, all stylet neurons, including these “Integrator” neurons, are not activated by high concentrations of nectar-specific sugars. Since glucose is a redundant cue in blood and nectar, coincident detection of multiple blood components in Integrator neurons confers context-specific information about glucose. These experiments reveal that upon initial contact with blood, specialized sensory neurons in the mosquito stylet innately encode the distinction between blood and nectar.

## RESULTS

### Sensory detection prior to blood- and nectar-feeding

Although both blood and nectar are appetizing to the female mosquito, blood-feeding and nectar-feeding are distinct feeding programs that utilize different peripheral sensory organs. When a female bites a human, she retracts the labium, uncovering the needle-like stylet required to draw blood. During blood feeding, the stylet pierces through skin to come into direct contact with blood while the labium remains on the surface of the skin (Figure 1A,B). During nectar-feeding, the labium directly contacts the nectar source and the stylet remains recessed and ensheathed within the labium (Figure 1C,D). In this configuration the stylet serves only as a feeding tube. Once feeding is initiated, there is a striking difference in the volume consumed and how these meals are metabolized by the digestive system after ingestion. The average blood meal is more than double the average sugar meal (Figure 1E,F) and the blood meal is immediately directed to the midgut for blood protein digestion, whereas the sugar meal is first directed to the crop (Figure 1G).

**Figure 1:**
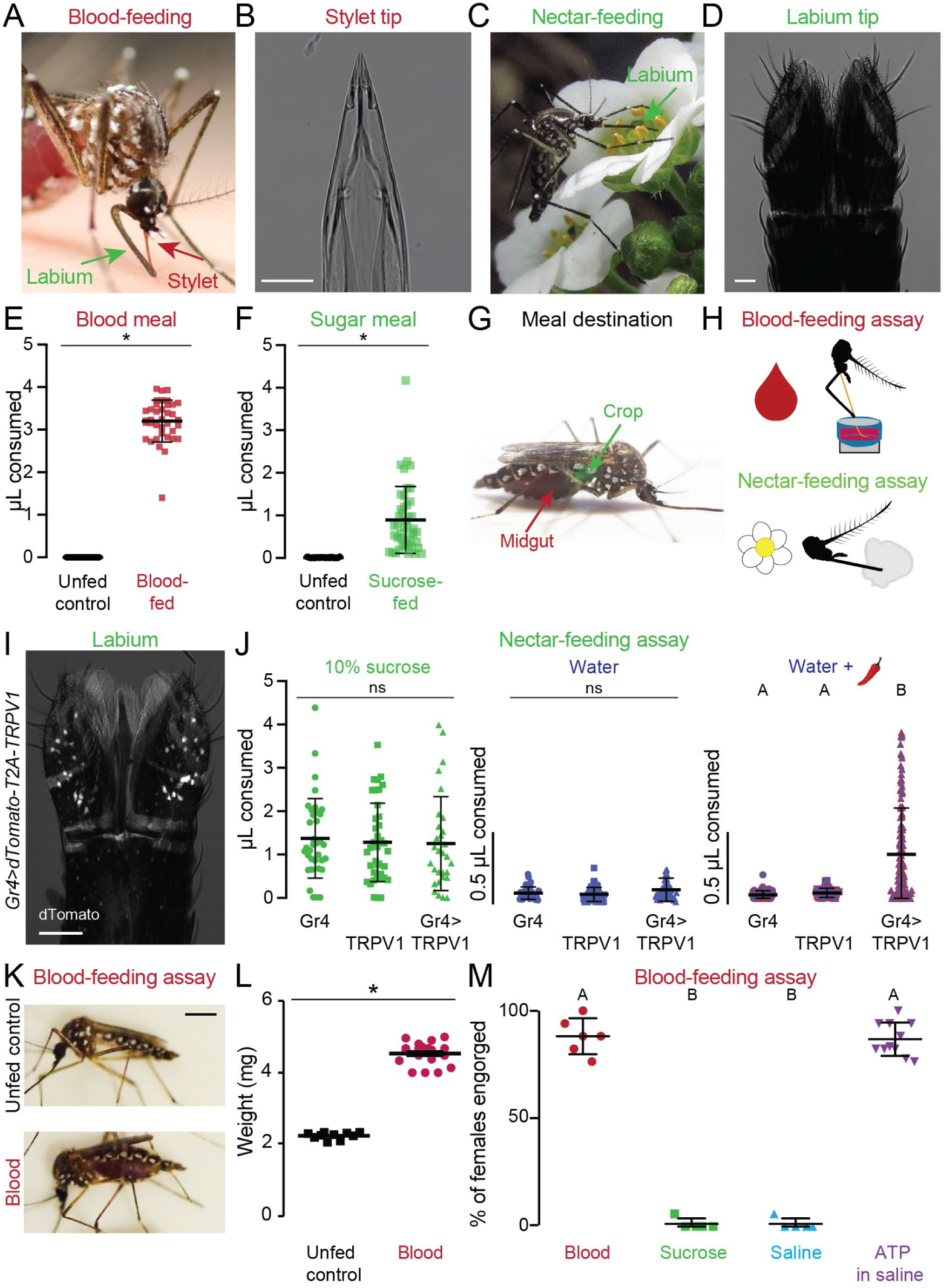
Sensory detection prior to blood- and nectar-feeding. (A,C) An *Ae. aegypti* female feeding on human skin (A, photo by Benjamin Matthews) or flower nectar (C, photo by Eric Eaton). (B,D) Transmitted light image of the female stylet (B) or labium (D). Scale bars: 25 µm (E,F) Volume of meal consumed after presenting blood (E) or sugar (F). Unfed controls were not given the option to feed and therefore represent the baseline for the assay. Each data point represents 1 female (mean±SD, N=37-46; * p < 0.05 Mann-Whitney test). (G) *Ae. aegypti* female with a blood meal in the midgut (red) and a 10% sucrose meal in the crop (green). Green food dye added to 10% sucrose to visualize meal location. (H) Schematic of blood- (top) and nectar-feeding (bottom) behavior assay. (I) Confocal image of dTomato expression in *Gr4>dTomato-T2A-TRPV1* labium with transmitted light overlay. Scale bar: 50 µm. (J) Volume of meal consumed by the indicated genotypes. Each data point represents 1 female: 10% sucrose N=30-40 females/genotype; water N=41-60 females/genotype; water + 50 µM capsaicin (red chili pepper): *Gr4* N=61, *TRPV1* N=62, *Gr4>TRPV1* N=124 females. (K) Female mosquitoes following 15 min exposure to different meals. Scale bar, 1 cm. (L) Sampled weight measurements from data for engorged females offered blood or unfed controls not offered any meal; N=10-19 weight measurements/meal (mean±SEM; * p < 0.05 unpaired t-test). (M) Female engorgement on the indicated meal delivered via Glytube. Each data point denotes 1 trial with 15-20 females/trial: N=5-11 trials/meal. In (J, M) data labeled with different letters are significantly different from each other (mean±SD; Kruskal-Wallis test with Dunn’s multiple comparison, p < 0.05).

Using standard laboratory blood-feeding or nectar-feeding assays (Figure 1H) (Costa-da-Silva et al., 2013; Liesch et al., 2013), we quantified features of each behavioral program and substituted different meal components to determine the requirements for feeding initiation. These assays mimic the circumstances of natural mosquito feeding behaviors. The blood-feeding assay offers females warmed meals in the presence of CO_2_ and heat, which attracts them to the artificial feeder (Liu and Vosshall, 2019; McMeniman et al., 2014). Upon landing, a parafilm membrane on the feeder separates the female from the meal, forcing her to pierce it with the stylet in the same way that she pierces skin to contact blood. In contrast, the nectar-feeding assay offers females room temperature meals on a cotton ball, allowing the labium to directly contact the meal upon landing.

To understand how mosquito nectar-feeding is initiated, we searched for orthologues of Gustatory Receptor (GR) genes that have been shown in *Drosophila melanogaster* flies to be selectively tuned to sweet tastants (Clyne et al., 2000; Dahanukar et al., 2001; Scott, 2018; Scott et al., 2001; Slone et al., 2007; Thorne et al., 2004). We identified *Ae. Aegypti Gr4* as the closest orthologue of both *Drosophila melanogaster Gr5a* and *Gr64f* (Kent and Robertson, 2009; Matthews et al., 2018). With the goal of labeling and manipulating neurons that express *Gr4*, we used CRISPR-Cas9 genome editing (Kistler et al., 2015; Matthews et al., 2019) to insert the QF2 transcriptional activator (Kistler et al., 2015; Matthews et al., 2019; Potter et al., 2010) at the endogenous *Gr4* locus. We also generated an effector QUAS line to express both the dTomato fluorescent reporter and the rat cation channel TRPV1 in *Gr4*-expressing neurons (Tobin et al., 2002). In *Gr4*>*dTomato-T2A-TRPV1* mosquitoes, we detected dTomato expression in the labium and legs, the two major taste appendages of insects (Figure 1I, Figure S1A-C). In our nectar-feeding assay, both the labium and leg can directly contact the meal during feeding, but the labium is the mouthpart used when feeding.

To ask whether activation of *Gr4* neurons is sufficient to initiate nectar-feeding behavior, we performed chemogenetic experiments that used capsaicin, the active ingredient in chili peppers, to activate TRPV1. Capsaicin should not affect feeding behavior of wild-type animals because capsaicin-sensitive TRP channels have not been described in invertebrates (Marella et al., 2006). In control experiments, we confirmed that capsaicin did not alter ingestion of water or sucrose by wild-type animals in the nectar-feeding assay (Figure S1D). Similar to previous observations in *Drosophila melanogaster* (Marella et al., 2006), addition of 50 µM capsaicin to water promoted ingestion of the otherwise inert water meal only in animals expressing TRPV1 in *Gr4* neurons (Figure 1J). Thus, nectar-feeding can be initiated by activation of sensory neurons that express sweet taste receptors.

What are the minimal sensory inputs required to initiate blood-feeding? When we used the blood-feeding assay to offer females warm sheep blood in the presence of heat and CO_2_, they reliably engorged on the meal, roughly doubling their initial body weight (Figure 1K-M). To separate meal composition from human cues, we maintained CO_2_ and heat delivery and exchanged the warm blood meal for warm sucrose or a saline solution that was isotonic with blood. Females consistently rejected both sucrose and saline in the blood-feeding assay, indicating that engorgement requires a separate step of evaluation after the female encounters a meal in the presence of human cues (Figure 1M). We then asked which blood meal components are important to trigger engorgement. Classic work from Hosoi and Galun indicated that the nutritional value of blood as a protein source can be uncoupled from blood-feeding behavior. Accordingly, an artificial meal consisting of blood proteins that is sufficient for egg production (Kogan, 1990) did not trigger engorgement unless ATP was added (Figure S1E,F). As previously reported, a protein-free solution of saline and ATP, or its non-hydrolyzable analogues, is sufficient for engorgement. (Figure 1M, Figure S1G,H) (Galun et al., 1963; Galun et al., 1984). This confirms that energy release from ATP hydrolysis can also be uncoupled from blood-feeding behavior. Finally, altering the concentration of ATP altered the probability of initiating engorgement (Figure S1I), but did not affect the meal size (Figure S1J). Together these behavioral data suggest that females can accurately recognize specific sensory features of blood and nectar to choose the appropriate feeding response.

### The stylet is poised to evaluate meal quality prior to blood-feeding

To understand how the taste of blood is recognized prior to blood-feeding, we first examined the stylet because it is the only sensory appendage to directly contact blood. We reasoned that if the stylet is assessing meal composition prior to engorgement, it must directly contact the meal both in situations where the mosquito decides to engorge and those where she does not. The blood-feeding assay gives a sensitive end-point measure of ingestion behavior but does not provide information about how and whether the stylet contacts the meal. To track the stylet of individual females in response to different meals presented with heat and CO_2_, we used the biteOscope assay (Hol et al., 2020). The biteOscope allowed us to visualize the stylet as it pierces a membrane and to determine whether the female subsequently engorged on warmed meals of water, saline, or ATP in saline (Figure 2A and Video 1). We selected ATP in saline as a proxy for blood since biteOscope meals must be optically clear to enable stylet video tracking. In all three conditions, the females repeatedly landed on the membrane and pierced it, bringing the stylet into direct contact with the meal, but females eventually engorged only on ATP in saline (Figure 2B-E). We conclude that human cues like heat and CO_2_ are sufficient to cause the female to pierce with her stylet and contact the meal, but additional blood-specific cues from the meal itself are required to trigger and sustain engorgement.

**Figure 2:**
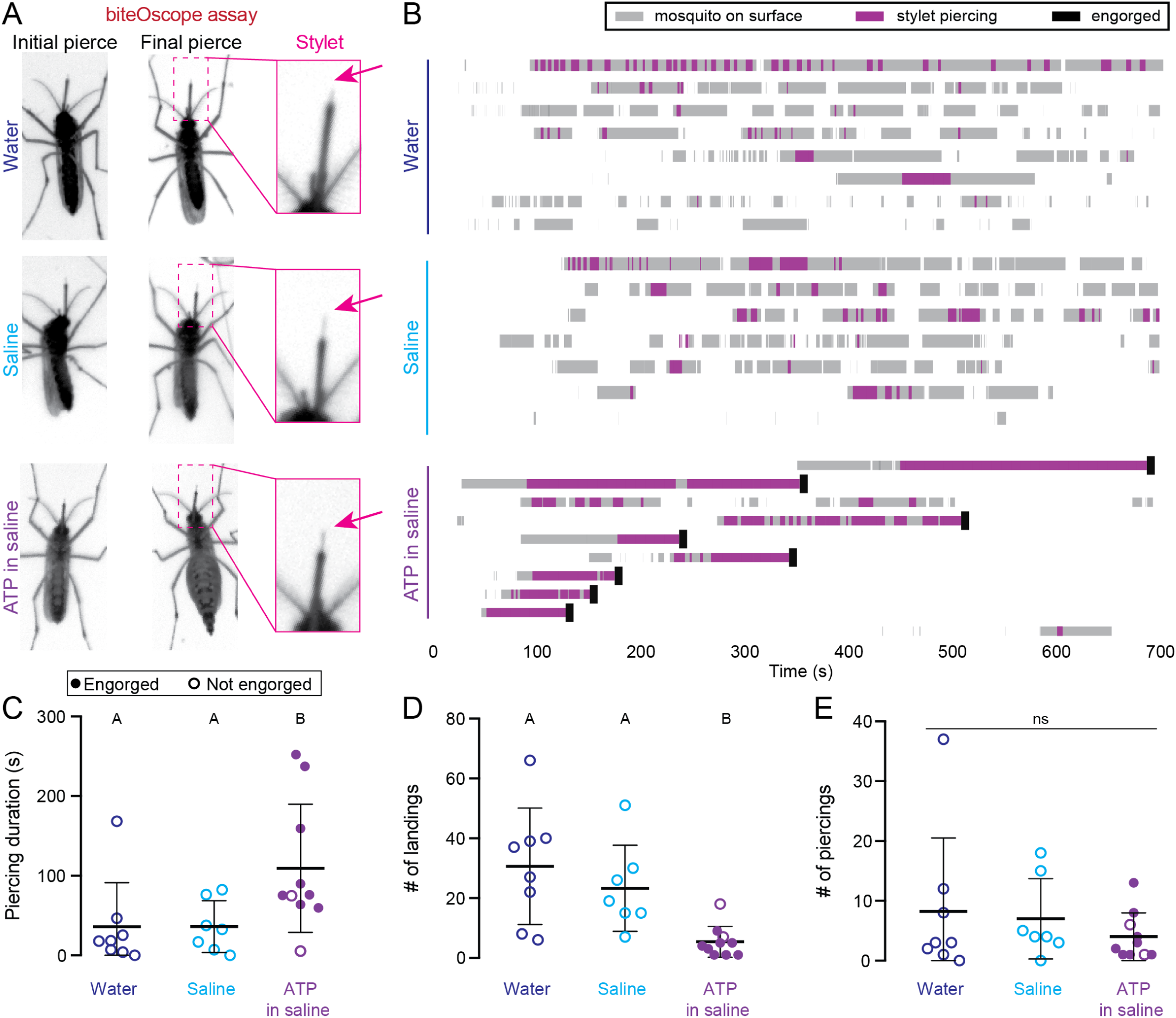
The stylet is poised to evaluate meal quality prior to blood-feeding. (A) Still video frames of female in biteOscope assay when stylet contacted meal for the first (left panel) or last (middle panel) time during the trial. Inset at right is from middle panel (see also Video 1). (B) biteOscope ethogram of landing events (gray boxes), stylet piercing events (pink boxes), and engorgement events (black boxes) for individual females provided water (N=8 females), saline (N=7 females), or 1mM ATP in saline (N=10 females) over 700 sec trial. Each row is an ethogram from 1 female. (C-E) Summary statistics from individual female ethograms in (A) for cumulative piercing duration during trial (C), # of landings (D), and # of piercings (E) for indicated meal. Each dot denotes 1 female, filled dot represents an engorged female. In C-E, data labeled with different letters are significantly different from each other (mean±SD; Kruskal-Wallis test with Dunn’s multiple comparison, p < 0.05).

### The stylet is sexually dimorphi

Since only female mosquitoes feed on blood, we hypothesized that a comparison of the female and male stylet would reveal the specialized sensory neurons involved in blood-feeding. Previous electron microscopy studies showed that females have exactly two sets of sensory sensilla, both of which are likely to directly contact blood underneath the skin (Lee, 1974). The first is a bilaterally symmetric set of two putative chemosensory sensilla, located at the distal tip and found only in the female stylet (Lee, 1974) (Figure S2A, pink arrows). The second is a bilaterally symmetric mechanosensory sensilla, located approximately 60 µm from the tip and found in both the female and male stylet (Lee, 1974) (Figure S2A, white arrows). Beyond this early description of the stylet sensillar morphology, there has been limited investigation of its neuroanatomy.

To reveal the organization of the stylet, we used reagents to stain cell nuclei and actin filaments, and visualized dTomato-labeled neurons in a *Brp*>*dTomato-T2A-GCaMP6s* pan-neuronal reporter strain (Figure 3A-F). Nuclear staining indicated that there is a concentration of rounder nuclei within the first 300 µm from the distal tip of the stylet, with more proximal nuclei showing a flatter elongated morphology (Figure S2B). When we examined dTomato expression in *Brp*>*dTomato-T2A-GCaMP6s* animals, we found that all stylet neurons are located within this distal region (Figure S2C). Moreover, we found that this section of the stylet is dramatically sexually dimorphic. When compared to males, females have a greater number of nuclei (Figure 3A,C), neurons (Figure 3B,D), and dendritic processes that innervate the distal tip (Figure 3E,F, Figure S2D-F). In agreement with previous electron microscopy data, we found mechanosensory sensilla innervation in both males and females (Figure 3E,F). These experiments illustrate the distinct neuroanatomy in the distal female stylet, consistent with blood-feeding being a female-specific behavior.

**Figure 3:**
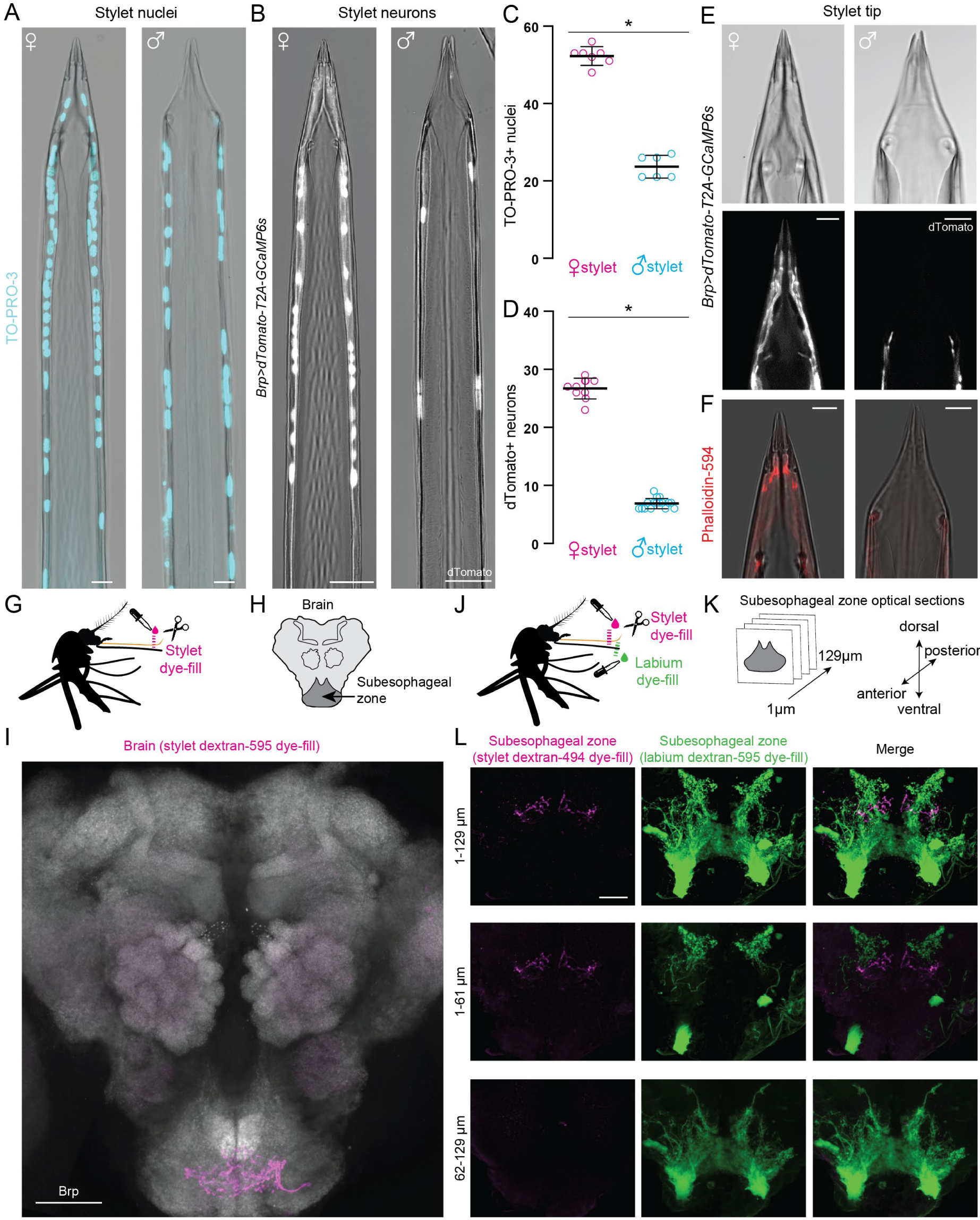
Sensory neurons in the female stylet are sexually dimorphic and project to a unique subesophageal zone region. (A,B) Confocal image with transmitted light overlay of TO-PRO-3 nuclear staining (cyan) in wild-type female (A, left) and male (A, right) stylets, and dTomato expression (gray) in *Brp>dTomato-T2A-GCaMP6s* female (B, left) and male (B, right) stylets. (C,D) Average # of TO-PRO-3 nuclei/stylet for most distal 300 µm (C, N=7 females, N=6 males), and dTomato neurons/stylet (D, N=10 females, N=16 males). Each dot denotes 1 animal (mean±SD, * p < 0.05 Mann-Whitney test). (E) Confocal image of transmitted light (top) and dTomato (gray, bottom) in *Brp>dTomato-T2A-GCaMP6s* female (left) and male (right) stylet tip. (F) Confocal image with transmitted light overlay of phalloidin-594 (red) staining in wild-type female (F, left) and male (F, right) stylets. (G, J) Schematic of stylet (G) and double (J) dye-fill experiment set-up performed in (I) and (L), respectively. (H, K) Schematic of mosquito brain region captured in (I), and subesophageal zone optical sections captured in (L). (I) Stylet neuron projection pattern (magenta) revealed by dextran-595 dye fill. Neuropil stained with anti-*Drosophila* Brp (gray). (L) Optical subesophageal zone sections from most anterior (top row) to most posterior (bottom row) of stylet (left, magenta) and labium (middle, green) projection pattern revealed by dual dextran-494 and dextran-595 dye-fill (see also Video 2). Scale bar: 25 µm (A,B,I,L), 10 µm (E,F).

We next asked where these female stylet neurons project in the mosquito brain. If the stylet detects the taste of blood, we would expect innervation of the subesophageal zone, the first processing center in the insect taste system (Ito et al., 2014; Scott, 2018). We performed dye-fill experiments to label axon terminals from all stylet neurons (Figure 3G) and found that stylet innervation was restricted to a discrete anterior and ventral region in the subesophageal zone (Ignell and Hansson, 2005) (Figure 3H, I). A previous study reported additional innervation of the antennal lobe, the primary olfactory processing center, upon dye-filling the stylet (Jung et al., 2015), but our results are inconsistent with this conclusion (Figure 3I and Figure S2G).

To determine if female stylet neurons and female labium neurons project to overlapping regions in the subesophageal zone, we performed a dual dye-fill experiment in which we labelled stylet and labium neurons with different dye colors in the same animal (Figure 3J). We found that the female labium projects to the posterior region of the subesophageal zone and that there is no direct overlap with stylet neuron projections (Figure 3K-L, Figure S2H, and Video 2). Therefore, inputs from the stylet and labium are segregated at the first synapse in the subesophageal zone.

### Stylet neurons detect blood

Our behavioral and anatomical results strongly suggest that stylet neurons can directly detect blood. We tested this by developing an *ex vivo* calcium imaging preparation with the pan-neuronal *Brp*>*dTomato-T2A-GCaMP6s* mosquito, which expresses both a dTomato marker and the genetically-encoded calcium indicator GCaMP6s (Chen et al., 2013) in all stylet neurons (Figure 4A,B). Because the stylet is a flat structure with all neurons located in one plane, we were able to image responses from all neurons simultaneously. When we applied 500 mM potassium chloride (KCl) as a depolarizing stimulus, we observed strong responses in all stylet neurons (Figure 4C). Since whole blood is opaque, it was necessary to devise a stimulation protocol that restricted blood to the stylet tip so that it did not interfere with GCaMP6s signal in the cell bodies. To solve this problem, we used the BioPen microfluidic device to deliver blood to the chemosensory pores that are innervated by sexually dimorphic distal processes (Figure 4D and Video 3). Stylet neurons consistently responded to 3 presentations of blood (denoted as 1^st^, 2^nd^, and 3^rd^ blood) separated by 60 sec intervals, and not to water (Figure 4E-I and Video 4). Within a given female, the peak ΔF/F response to multiple presentations of blood was stable, but the exact number and position of blood-sensitive neurons was not stereotyped across individuals (Figure 4G,H). We also noted that different neurons within an individual had unique GCaMP6s response waveforms and that these waveforms were stable across every blood presentation for a given neuron (Figure 4I). Across individuals we observed that approximately 50% of stylet neurons responded to blood (Figure 4J). These results demonstrate that a large population of stylet chemosensory neurons responds directly to whole blood.

**Figure 4:**
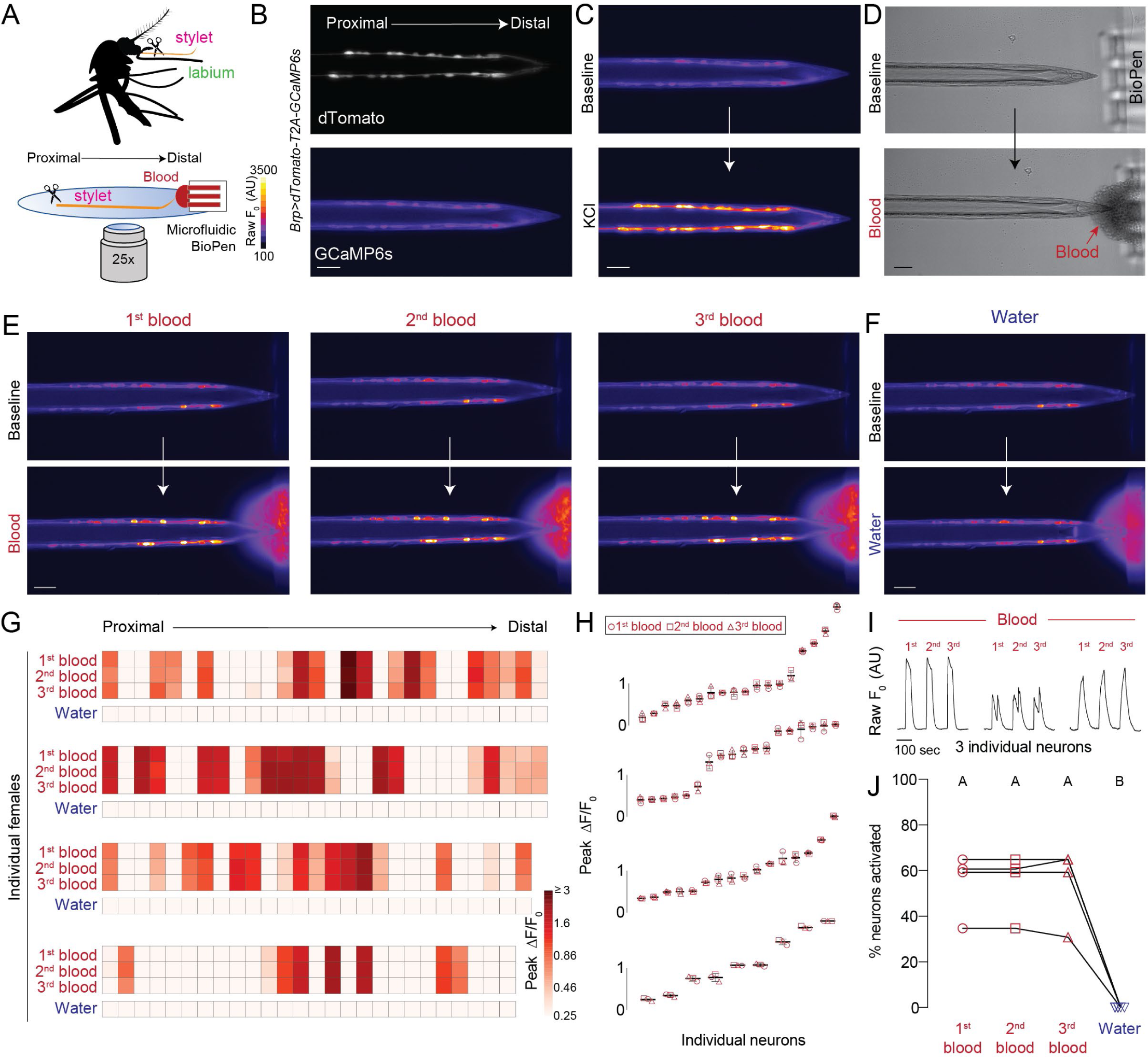
Sexually dimorphic stylet neurons directly sense blood. (A) Schematic of *ex vivo* stylet imaging preparation. (B) Wide-field image of dTomato (top) and baseline GCaMP6s (bottom, scale: arbitrary units) for a representative stylet, oriented proximal to distal. (C) Representative image of GCaMP6s fluorescence increase to bulk neuronal depolarization with 500 mM KCl (bottom) compared to base-line (top). (D) Representative bright-field image before (top) and during (bottom) delivery of sheep blood to the stylet tip via the BioPen (see also Video 3). (E,F) Representative image of GCaMP6s fluorescence increase to indicated blood presentation (bottom, E) or water control (bottom, F), compared to baseline (top) (see also Video 4) (G) Heat maps of peak ΔF/F_0_ response to the indicated ligand. Each square is the average of 3 ligand exposures and each column represents one neuron. Each row represents the response to indicated ligand for all neurons from 1 individual female, with neurons ordered from proximal to distal. N=4 individual females. (H) Summary of peak ΔF/F_0_ data in (G) for neurons with ≥ 0.25 peak ΔF/F_0_ to at least 1 of 3 blood presentations. Each data point represents an average of 3 ligand exposures (mean±SD) and are sorted by peak ΔF/F_0_ (I) A subset of traces for 3 neurons from 1 individual in (H), y axis scale: arbitrary units of raw fluorescence. (J) Quantification from (G, H) of % neurons with ≥ 0.25 peak ΔF/F_0_ to the indicated ligand, each data point denotes the response from 1 female and responses from the same female are connected by a line. Data labeled with different letters are significantly different from each other (one-way repeated measures ANOVA, with the Geisser-Greenhouse correction and Tukey’s multiple comparisons test, p < 0.05). (B-F) Scale bar: 25 µm. 0.0002% fluorescein was added to blood and water stimuli to visualize ligand delivery zone.

### Blood detection is combinatorial across taste qualities

How is blood, a complex mixture of cells, proteins, lipids, metabolites, and salts, represented by stylet neurons? We used a reductionist approach to understand how the taste of blood is encoded in stylet neurons. We selected 4 blood components [adenosine triphosphate (ATP), glucose, sodium bicarbonate (NaHCO_3_), and sodium chloride (NaCl)] that have been individually shown to increase the probability of engorgement (Galun et al., 1984; Gonzales et al., 2018). ATP and unbuffered NaHCO_3_ (pH = 8 - 9) are not associated with canonical taste qualities, but glucose and sodium chloride are traditionally associated with sweet and salty, respectively. We selected concentrations of glucose, sodium bicarbonate, and sodium chloride within range of standard blood values for vertebrate species. For ATP, it is difficult to determine the exact *in vivo* concentration present when the female bites a human because ATP is derived from multiple sources and is rapidly hydrolyzed. Up to millimolar-range ATP can be released from the deformation or lysis of red blood cells or from epithelial cells lining the blood vessel as a damage response to the stylet piercing (Born and Kratzer, 1984; Forsyth et al., 2011). At steady-state, free ATP in plasma is present in the nanomolar-range (Gorman et al., 2007). We selected 1 mM because it resulted in the most robust responses in the behavioral dose response curve (Figure S1I). Using the blood-feeding assay, we found that the combination of these 4 ligands (hereafter referred to as Mix+ATP) was sufficient to reliably trigger engorgement (Figure 5A,B).

**Figure 5:**
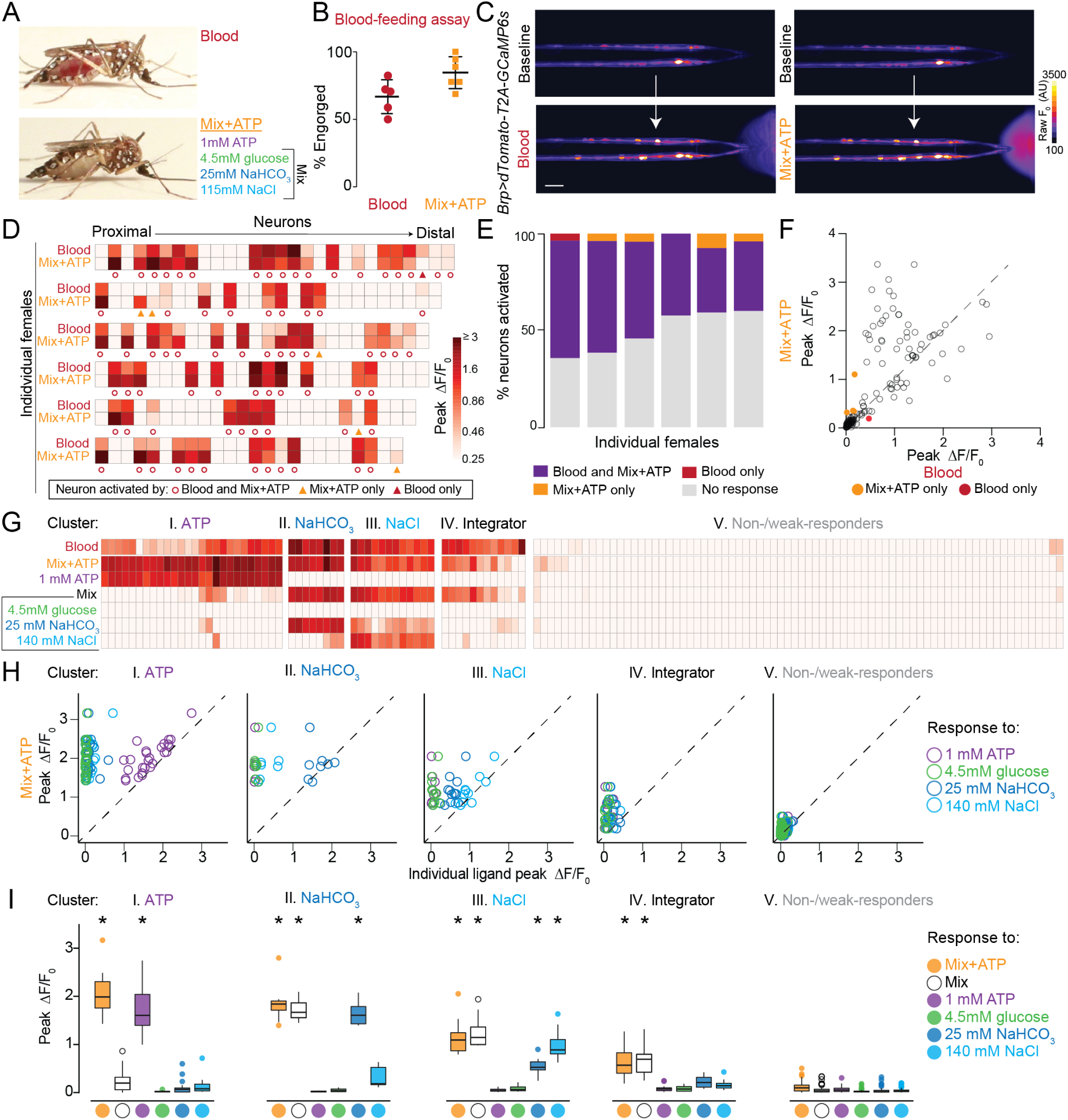
Stylet neurons integrate across taste modalities to detect blood. (A) Representative engorged *Ae. aegypti* female following 15-min exposure to blood (top) or Mix+ATP (bottom) via Glytube assay. (B) Female engorgement on blood (N=5 trials) and Mix+ATP (N=6 trials) delivered via Glytube (lines denote mean±SD, 15–20 females/trial, one-sample t-test relative to 0, p = 0.0003, p < 0.0001). (C) Representative image of GCaMP6s fluorescence increase (scale: arbitrary units) to blood (bottom, left) or Mix+ATP (bottom, right), compared to baseline (top) (See also Video 5). Scale bar: 25 µm. (D) Heat maps of peak ΔF/F_0_ response to the indicated ligand. Each square is the average of 3 ligand exposures and each column represents one neuron. Each row represents the response to indicated ligand for all neurons from 1 individual female, with neurons ordered from proximal to distal. N=6 individual females. (E) Summary of % neurons with ≥ 0.25 peak ΔF/F_0_ to the indicated ligand from (D), each column represents 1 female. (F) Scatter plot comparing peak ΔF/F_0_ in response to Mix+ATP (y-axis) and blood (x-axis) summarized across N=6 females from (D,E). Each dot represents 1 neuron, dots that fall on the dashed line have the same peak ΔF/F_0_ in response to blood and Mix+ATP. (G-I) 5 clusters of blood-sensitive neurons identified by unsupervised hierarchical clustering of peak ΔF/F responses to the ligands indicated in (G). Clustering removes proximal-distal ordering and female identity. N=5 females (* p < 0.05, one-sample Wilcoxon signed-rank test). (A-I) In this and all subsequent experiments “Mix” is 4.5 mM glucose, 25 mM NaHCO_3_, 115 mM NaCl and “Mix+ATP” is Mix supplemented with 1 mM ATP. To visualize ligand delivery zone, 0.0002% and 0.00002% fluorescein was added to blood and Mix+ATP, respectively, in BioPen experiments.

Since both blood and Mix+ATP trigger engorgement, are there differences in how stylet neurons respond to these taste stimuli? When we delivered blood or Mix+ATP to *Brp*>*dTomato-T2A-GCaMP6s* animals, we found that blood and Mix+ATP activate the same population of stylet neurons (Figure 5C-F and Video 5). Although the magnitude of response can vary within a given neuron (Figure 5D,F), we consistently saw Mix+ATP-responsive neurons tracking with blood-responsive neurons across individuals, irrespective of variability in the position of the neuronal cell body along the proximal-distal axis of the stylet (Figure 5D,E).

To understand how blood components contribute to the perception of whole blood, we used Mix+ATP as a chemically-defined mixture that reliably activates blood-responsive neurons. When we presented each component of Mix+ATP individually, we found that blood-sensitive neurons are a heterogenous population and that different neuronal subsets within each female can respond to distinct blood components (Figure S3A,B). Moreover, all components except 4.5 mM glucose reliably activated subpopulations of stylet neurons when presented individually. Unsupervised hierarchical clustering of this dataset was performed to group neurons into 5 clusters according to their functional response profile (Figure 5G). For each neuron in a cluster, we calculated a ratio of peak ΔF/F response to Mix+ATP compared to the peak ΔF/F response to any individual ligand (Figure 5H). We found that the first 3 clusters represent neurons reliably activated by an individual component: ATP, sodium bicarbonate, and sodium chloride, respectively (Figure 5I). Although Cluster 4 was not reliably activated by any individual ligand, it was reliably activated by a mixture of sodium bicarbonate, sodium chloride, and glucose (hereafter referred to as “Mix”) (Figure 5I). We define these as “Integrator” neurons and explore their function in subsequent experiments. Cluster 5 neurons were non-responsive or showed weak responses (Figure 5I). Together these experiments demonstrate that subsets of blood-sensitive neurons are selectively tuned to specific blood components that span multiple canonical and noncanonical taste qualities.

### *Ir7a* and *Ir7f* mark functionally distinct populations of blood-sensitive neurons

We next asked if these functionally distinct blood-sensitive subsets are transcriptionally-defined populations. To identify candidate genetic markers for neuronal subsets, we searched for transcripts expressed only in the female stylet and not in tissues that do not directly contact blood. We profiled gene expression in the female stylet using RNA-seq and compared it to the male stylet and the female labium. To identify female stylet-specific transcripts, we analyzed the intersection of genes significantly enriched in the female stylet compared to both the female labium and the male stylet (Figure S4A,B, fuchsia data points). We further filtered for transcripts that were expressed at very low levels (< 0.5 transcripts per million, TPM) in a comprehensive transcriptome dataset that included other sensory appendages, brain, and ovary (Figure S4C) (Matthews et al., 2018; Matthews et al., 2016). Of the four transcripts that met these criteria for female stylet-specific expression, two were members of the ionotropic receptor (IR) superfamily, *Ir7a* and *Ir7f* (Figure S4D). Since IRs have been shown to play roles in chemo-, thermo-, and mechano-reception (Benton et al., 2009; Rytz et al., 2013), we reasoned that *Ir7a* and *Ir7f* were likely to be expressed in sensory neurons.

To identify and characterize the neurons that express these female stylet-specific transcripts, we generated QF2 driver lines for *Ir7a* and *Ir7f*. When crossed to reporter lines, both showed sparse expression in subsets of chemosensory neurons in the female stylet (Figure 6A,B). *Ir7a* and *Ir7f* are expressed in approximately 1 - 2 neurons and 3 - 4 neurons, respectively, in female stylets. No expression of either gene was detected in male stylets. The sparse nature of these drivers allowed us to more carefully examine dendritic innervation of the bilaterally symmetric set of two chemosensory sensilla at the stylet tip. For each neuron in *Ir7a* animals, we found ipsilateral innervation of the proximal sensilla (Figure 6A). For each side of the stylet in *Ir7f* animals, the proximal neuron ipsilaterally innervated the proximal sensilla, and the distal neuron ipsilaterally innervated the distal sensilla (Figure 6B). When we examined the projection patterns of these neurons in the female brain, we observed axons innervating the same ventral subesophageal zone region identified in our stylet dye fills (Figure 6C-F). Importantly, no regions in the male brain or additional regions in the female brain were labeled in these strains, highlighting the exquisite selectivity of *Ir7a* and *Ir7f* gene expression to the female stylet. We used these markers to study the properties of these neuronal populations, rather than the properties of the genes themselves.

**Figure 6:**
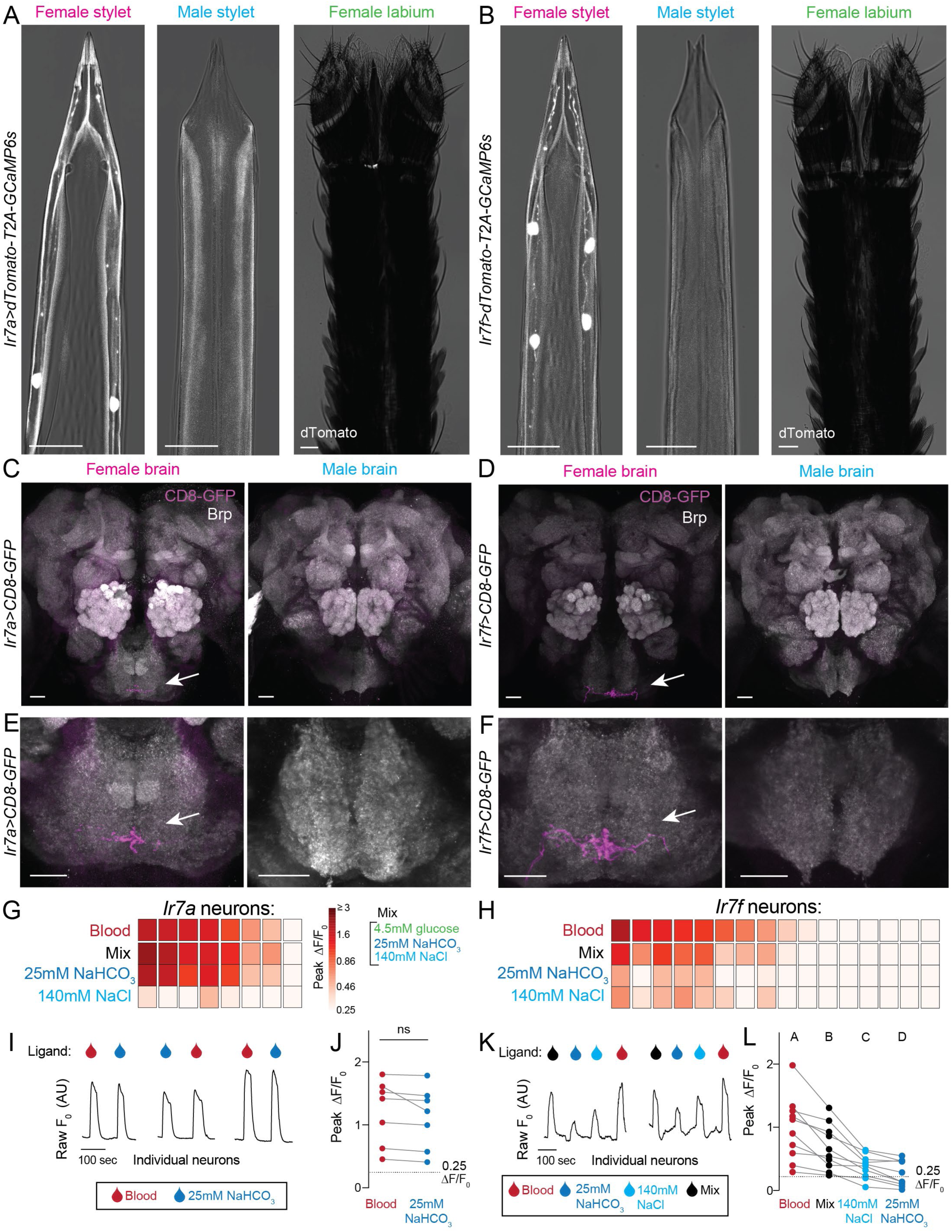
*Ir7a* and *Ir7f* mark the sodium bicarbonate and Integrator neurons. (A,B) Confocal image with transmitted light overlay of dTomato expression (gray) in the female stylet (left panel), male stylet (middle panel), and female labium (right panel) of *Ir7a*>*dTomato-T2A-GCaMP6s* (A) and *Ir7f*>*dTomato-T2A-GCaMP6s* (B) animals. *Ir7a* expression: 10/13 females = 2 neurons, 2/13 females = 1 neuron, 1/13 females = 0 neurons. *Ir7f* expression: 6/11 females = 4 neurons, 5/11 females = 3 neurons. (C-F) mCD8:GFP expression (magenta, white arrow) of *Ir7a*>*mCD8:GFP* (C,E) and *Ir7f*>*mCD8:GFP* (D,F) in female (left) and male (right) brain (top) and subesophageal zone (bottom). Neuropil labeled with anti-*Drosophila* Brp (gray). Brain and subesophageal zone images were acquired from two different individuals. (G,H) Heat maps of peak ΔF/F_0_ response to the indicated ligand in *Ir7a*>*dTomato-T2A-GCaMP6s* (G) and *Ir7f*>*dTomato-T2A-GCaMP6s* (H) neurons across N=5 females. Each square is the average of 3 ligand exposures and each column represents one neuron. Columns are sorted by largest to smallest peak ΔF/F_0_ in response to blood. (I,K) Raw F_0_ traces from individual neurons in response to indicated ligand. (J,L) For blood-sensitive neurons, peak ΔF/F_0_ to indicated ligand. Each data point denotes the response from 1 neuron and responses from the same neuron are connected by a line. In (J), ns (not significant), p > 0.05, paired t-test. Data in (L) labeled with different letters are significantly different from each other (one-way repeated measure ANOVA, with the Geisser-Greenhouse correction and Tukey’s multiple comparisons test, p < 0.05). Scale bar: 25 µm. 0.0002% fluorescein was added to blood and 140 mM NaCl, and 0.00002% was added to Mix or 25 mM sodium bicarbonate in the BioPen to visualize ligand delivery zone.

To determine the functional properties of *Ir7a* and *Ir7f* neurons, we performed cell-type specific calcium imaging experiments and found that across the animals tested, almost all *Ir7a* neurons and approximately half of *Ir7f* neurons respond to blood (Figure 6G,H). We then tested responses to the individual components of Mix+ATP and found that both blood-responsive populations respond to Mix (glucose, sodium bicarbonate, and sodium chloride) and not ATP. *Ir7a* blood-sensitive neurons are most robustly activated by sodium bicarbonate (Figure 6I,J), sharing a profile with sodium bicarbonate neurons identified in Cluster II (Figure 5G-I). In contrast, *Ir7f* blood-sensitive neurons were activated robustly by Mix and had moderate and variable responses to 140 mM sodium chloride and/or 25 mM sodium bicarbonate (Figure 6K,L), sharing a profile most similar to Integrator neurons in Cluster IV (Figure 5G-I). Thus these two female stylet-specific driver lines define the molecular and functional identity of two non-overlapping blood-sensitive neuron populations in the female stylet.

### Specialization in stylet neurons enables discrimination between blood and nectar

To avoid confusion between the blood-feeding and nectar-feeding programs, it is important that a female not trigger engorgement when the stylet contacts nectar sugars during blood-feeding. How is this discrimination achieved? To address this question, we investigated behavioral and neuronal responses to nectar sugars.

Sugars present an interesting discrimination challenge for mosquito taste coding because a female wants to recognize nectar as appetizing when she intends to feed on nectar, but not when she intends to feed on blood. To further complicate matters for the mosquito, glucose is a redundant cue in blood and nectar (Figure 7A). Since stylet neurons are the only sensory neurons that directly contact the meal during blood feeding, do they have a specialized taste coding strategy to selectively distinguish blood components from nectar components? One way to achieve this discrimination is to segregate the expression of classical sweet taste receptors in the labium and prevent their expression in the stylet. Alternatively, stylet neurons could express the same sweet taste receptors that mediate nectar detection in the labium, but sweet taste sensitivity and/or processing could be rapidly modulated during blood-feeding by the presence of human cues like CO_2_ and heat. If the stylet has a canonical sweet taste pathway, we would expect the stylet to express sweet gustatory receptors and respond to sucrose, fructose, and glucose, which are the principal components of nectar (Liman et al., 2014; Yarmolinsky et al., 2009).

**Figure 7:**
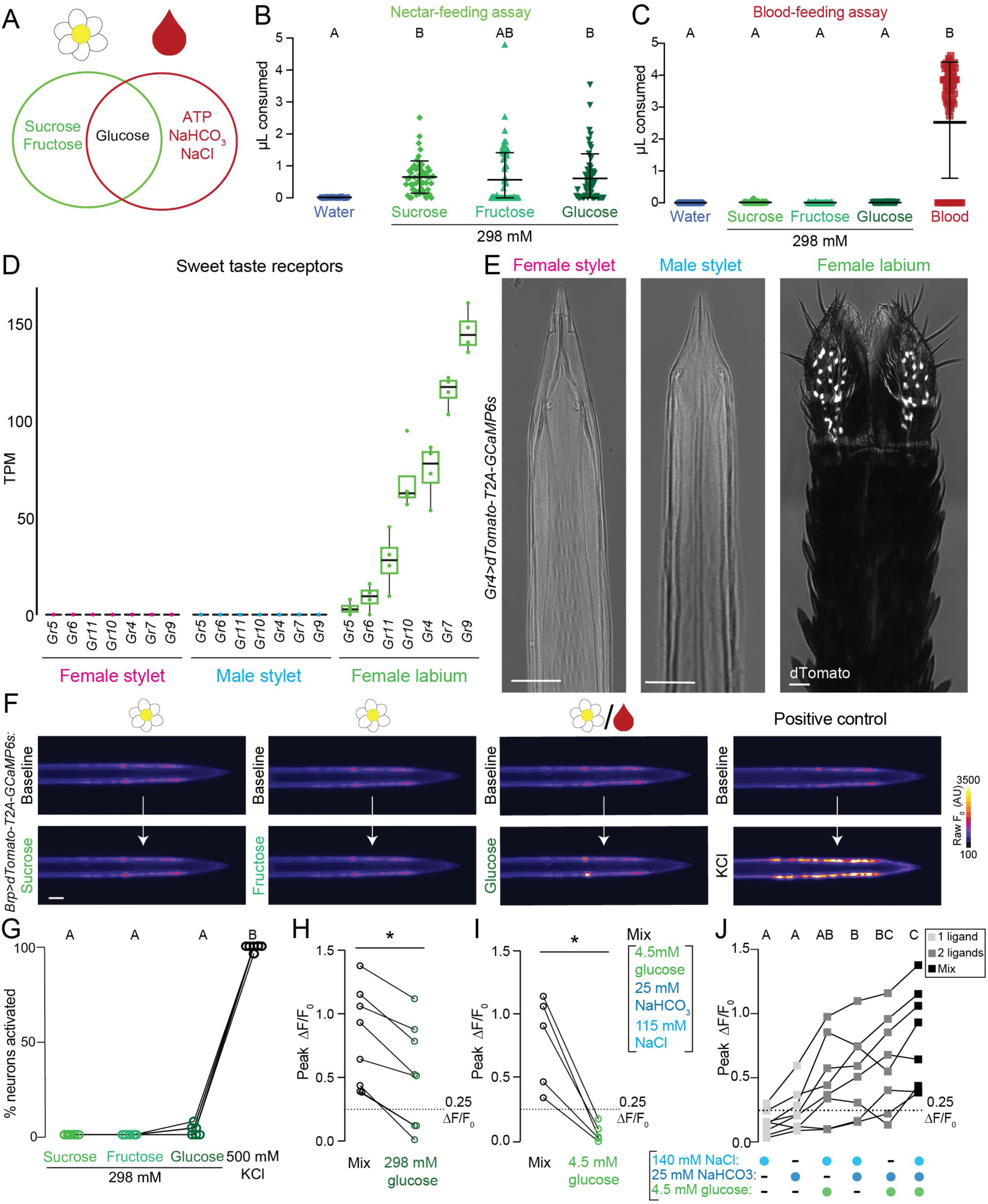
The stylet is specialized to detect blood over nectar. (A) Venn diagram schematizing the similarity and differences between nectar (left circle) and blood (right circle) components. (B,C) Volume of indicated meal consumed in the nectar-feeding (B) and blood-feeding (C) assay. Each data point represents 1 female: water N=36-40; sucrose N=53–60; fructose N=40-74; glucose N=55-59. Blood in (C) is a positive control for blood-feeding assay, N=76 females. (D) Sweet taste receptor expression from RNA-seq analysis of the indicated tissues. N=4 replicates/tissue. Median indicated by black line, bounds of box represent first and third quartile, whiskers are 1.5 times the inter-quartile range and outliers denoted by dot. (E) Confocal image with transmitted light overlay of dTomato expression (gray) in the female stylet (left panel), male stylet (middle panel), and female labium (right panel) of *Gr4*>*dTomato-T2A-GCaMP6s* animals. (F) Representative image of GCaMP6s fluorescence increase to indicated 298 mM sugar presentation (bottom) compared to baseline (top). Flower/blood symbol indicate that sugar is found in nectar and blood. (G) Quantification of % neurons with ≥ 0.25 peak ΔF/F_0_ to the indicated ligand, each data point denotes the response from 1 female, responses from the same female are connected by a line, N=6 females.(H,I) For Integrator neurons, peak ΔF/F_0_ to 298 mM glucose (H, N=8 neurons) and 4.5 mM glucose (I, N=5 neurons). Each dot represents 1 neuron (mean±SD, * p < 0.05 Mann-Whitney test). (J) For Integrator neurons, peak ΔF/F_0_ to indicated ligand(s). Each data point denotes the response from 1 neuron, N=8 neurons. Data labeled with different letters are significantly different from each other (one-way repeated measures ANOVA, with the Geisser-Greenhouse correction and Tukey’s multiple comparisons test, p < 0.05). Scale bar: 25 µm. (B,C,G) Data labeled with different letters are significantly different from each other (mean±SD; Kruskal-Wallis test with Dunn’s multiple comparison, p < 0.05). (H, I, J) Responses from the same neuron are connected by a line.

We first measured the behavioral response to nectar sugars in the context of the nectar- and blood-feeding assay. We presented each nectar component at 298 mM because this concentration is approximately equivalent to the female’s normal sugar meal that is sufficient for energy metabolism (Van Handel, 1972, 1984). Females readily ingested all three sugars when the labium directly contacted the meal in the nectar-feeding assay, where no host cues are present (Figure 7B). In contrast, they rejected these same sugars in the blood-feeding assay when the stylet directly contacted the meal in the presence of heat and CO_2_ (Figure 7C). In control experiments we showed that blood stimulated robust consumption in the blood-feeding assay (Figure 7C).

These results lead to the question of whether stylet neurons can detect these nectar sugars at all. When we examined the RNA-seq dataset, we found the female labium expresses all predicted orthologues of *Drosophila melanogaster* canonical sugar receptors (Figure 7D). However, none of these sweet gustatory receptor transcripts were detected in the female or male stylet (Figure 7D). Confirming this observation, the *Gr4* reporter line showed no expression in the female and male stylet but is expressed in a large group of labial neurons (Figure 7E). Moreover, these labial neurons project axons to the posterior region of the subesophageal zone that we identified in our labium dye-fill experiments, but not to the anterior, ventral region occupied by stylet neuron projections (Figure S5A,B). Thus stylet neurons do not express canonical sweet taste gustatory receptors.

We next examined stylet neuron responses to nectar sugars using calcium imaging. If stylet neurons lack a canonical sweet taste pathway, we expect that they would not generically respond to sucrose, fructose, and glucose. To test this, we presented these sugars at 298 mM and measured neuronal responses in *Brp*>*dTomato-T2A-GCaMP6s*. To control for osmotic effects, stylets were pre-equilibrated in 298 mM of the behaviorally inert/non-sweet sugar cellobiose for 30 sec prior to the isomolar sugar of interest. Indeed, no stylet neurons responded to 298 mM nectar-specific sugars (Figure 7F,G). We observed occasional responses to 298 mM glucose, which is the only sugar found in both blood and nectar (Figure 7F,G). In positive control experiments, we confirmed that stylet neurons responded to 500 mM KCl (Figure 7F,G). Together these results demonstrate that the stylet has a unique taste coding strategy that is specialized to detect blood and blood components over nectar-specific components.

Although responses to 298 mM glucose were less frequent, we found that 298 mM glucose-sensitive neurons were blood sensitive and shared a functional profile with Integrator neurons (Figure S5C,D). Integrator neurons consistently responded more to Mix than this high concentration of glucose. (Figure 7H). We therefore asked if physiological levels of blood glucose directly contribute to Mix responses that we observed in Integrator neurons. Since Integrator neurons do not respond to 4.5 mM glucose alone (Figure 5G-I and Figure 7I), we tested if the addition of 4.5 mM glucose to other Mix components increases the total neuronal response. Integrator neurons respond reliably to 4.5 mM glucose when co-presented with sodium chloride or sodium bicarbonate and are maximally activated by co-presentation of all three (Figure 7J). These results demonstrate that individual sensory neurons can directly integrate glucose (sweet), sodium chloride (salty), and sodium bicarbonate. Polymodal integration in taste cells is an unexpected finding because these different taste qualities activate mutually exclusive taste cell populations in *Drosophila* and rodents (Liman et al., 2014; Yarmolinsky et al., 2009). We propose that in the mosquito, this unusual taste coding strategy enables flexible responses to physiological levels of blood glucose depending on the context in which it is detected. Since a female could encounter this concentration of glucose in blood or nectar, polymodal integration is an additional mechanism for the stylet to selectively recognize the taste of blood. Taken together, our results demonstrate that the stylet is specialized to detect blood over nectar.

## DISCUSSION

In this study we show that sexually dimorphic stylet neurons are the first sensory neurons to detect blood as an *Ae. aegypti* female draws a blood meal. Using pan-neuronal calcium imaging, we find that stylet neurons taste multiple blood components to form the percept of blood. We discovered that stylet neurons are specialized to detect blood over nectar, facilitating peripheral discrimination between these two appetizing food sources during blood-feeding.

### Anatomical, molecular, and functional properties of the stylet

The female stylet is an unconventional sensory organ whose functional properties are poorly understood. The microneedle-like biophysical properties needed to efficiently pierce skin (Choumet et al., 2012; Ramasubramanian et al., 2008) may influence its unique anatomical organization into two single-file rows of cells along each side. Consistent with blood-feeding being specific to females, we identified dramatic sexual dimorphism in neuron number and innervation of chemosensory sensilla. The sparse, stylet-specific *Ir7a* and *Ir7f* driver lines reveal that individual neurons send ipsilateral dendrites into one of the two chemosensory sensilla found on each side of the stylet tip. Interestingly, we observed inter-individual differences in blood-sensitive neuron number and cell body position. We do not yet understand the mechanism of developmental patterning that permits variable cell body position along the proximal-distal axis of the stylet. Variability in the exact distance of the cell to the stylet tip may be tolerated because all stylet neuron dendrites terminate at the tip, irrespective of cell body position.

By generating two female stylet-specific driver lines, we revealed non-overlapping blood-sensitive neurons belonging to two functionally distinct subsets: *Ir7f* blood mixture-sensitive neurons and *Ir7a* sodium bicarbonate-sensitive neurons. Together, these driver lines reveal the molecular identity of approximately one quarter of total stylet neurons. Future work will allow us to determine if *Ir7a* and *Ir7f* transcripts, along with additional putative chemosensory receptors identified in our stylet RNA-seq dataset, contribute to blood ligand detection. This dataset is also an important resource to help identify genetic markers for sodium chloride- and ATP-sensitive subsets and to resolve the complete molecular landscape of stylet neurons.

A major finding of this work is that four ligands previously shown to increase the probability of initiating blood-feeding behavior do indeed directly activate the stylet. When presented as a mixture, these four blood components—ATP, glucose, sodium bicarbonate, and sodium chloride—are sufficient to initiate blood-feeding behavior and activate the same neurons as blood. It is surprising that females will so readily engorge on Mix+ATP because it does not contain the proteins required for egg development. It may be that proteins are not an ideal substrate for blood recognition because their amino acid sequences and structures can evolve over time and across species. In contrast, the chemical structures of ATP, glucose, sodium bicarbonate, and sodium chloride are invariant across evolutionary time and species.

Our functional imaging shows that roughly half of the 40 stylet neurons can be activated by blood. The remaining stylet neurons may respond to a variety of different ligands, including ligands found only when the stylet contacts an intact capillary microenvironment. For example, once blood is drawn, the concentration of certain volatile or unstable blood components is likely to decrease. There may also be circulating factors released from surrounding cells as a damage response to the piercing stylet or ligands specific to human blood. These unidentified ligands may be detected in an *in vivo* context, but none appear to be required for blood-eeding behavior or egg development. Another possibility is at least some of the remaining stylet neurons respond to additional taste qualities observed in other feeding appendages. For example, responses to osmolarity, high salt, CO_2_, and bitters have been observed in labellar neurons in *Drosophila melanogaster* (Liman et al., 2014; Yarmolinsky et al., 2009). Bitters are of particular interest because specific bitters can be added to blood and prevent feeding (Dennis et al., 2019). Finally, the stylet could be capable of thermosensation or mechanosensation related to blood flow or tissue penetration. The pan-neuronal stylet imaging preparation we have developed will facilitate future systematic analyses of stylet responses to diverse sensory stimuli.

### Stylet neurons integrate across taste qualities to detect blood

We found that blood-sensitive neurons can be divided into functionally distinct subtypes, each identifiable by their neuronal response to the panel of ligands we tested. Each subset is activated by a behaviorally-relevant concentration of a ligand, or mix of ligands, found in blood. Glucose and sodium chloride are associated with the distinct taste qualities of sweet and salty, but it is not clear if sodium bicarbonate or ATP responses overlap with a canonical taste quality. In blood, sodium bicarbonate is buffered at pH = 7.4 and predominately present as HCO_3_-, with 10% or less present as CO_2_ (Centor, 1990). CO_2_ underlies sour taste and the taste of carbonation, but HCO_3_-has not yet been assigned to a defined taste quality. Similarly, there is no description of the taste of ATP. The closest comparison to ATP detection is in mammals, where specific 5’-monophosphate nucleotides potentiate umami perception (Yamaguchi, 1967) and in *Drosophila melanogaster* larvae, where certain ribonucleosides directly activate *Gr28*-expressing taste neurons (Mishra et al., 2018). Our work shows that the taste of blood is multidimensional and that multiple taste qualities, both canonical and noncanonical, are integrated across subsets of blood-sensitive neurons and for the particular subset of Integrator neurons, within individual neurons.

We directly observed integration by a subset of stylet neurons maximally activated by co-presentation of glucose, sodium chloride, and sodium bicarbonate. Simultaneous detection of sweet, salty, and sodium bicarbonate in one neuron is unexpected because distinct taste qualities are thought to activate non-overlapping sensory neuron populations in both mammals and insects (Yarmolinsky et al., 2009). Yet here we only detect responses to physiological levels of blood glucose (4.5 mM) in the presence of sodium chloride or sodium bicarbonate. We speculate that polymodal neurons act as coincidence detectors and that 4.5 mM glucose alone produces subthreshold responses without the co-presentation of sodium chloride and/or sodium bicarbonate. Since glucose is a redundant cue in blood and nectar, this unconventional taste coding mechanism confers an important distinction between glucose present in blood versus nectar.

What is the molecular mechanism of glucose detection and integration in polymodal stylet neurons, given that no canonical sweet taste receptors are detected in the stylet? It remains unknown if one receptor can directly integrate the chemically distinct ligands of sodium bicarbonate, glucose, and sodium chloride, or if the neuron integrates activity from multiple independent receptors. Although we found no canonical sweet gustatory receptor expression in the stylet, the stylet does express *Gr34*, an orthologue of *D. melanogaster Gr43a* (Matthews et al., 2018). *D. melanogaster Gr43a* does not share sequence homology with the sweet taste subfamily and is the only gustatory receptor also expressed in the brain, where it acts as a metabolic sensor of circulating fructose (Miyamoto et al., 2012). *Gr34* may be repurposed as a glucose receptor in the stylet or there may be an unconventional receptor for glucose in these neurons. Since *Ir7f*-expressing neurons intersect with the Integrator neuron subset, their molecular profile may help uncover this mechanism of taste quality integration.

Taste quality integration also occurs across the distinct blood-sensitive neuronal subsets to form the neural representation of blood in the stylet. We found that behaviorally-relevant concentrations of ATP, sodium bicarbonate, and sodium chloride were individually sufficient to activate a subset of stylet neurons. However, any individual component was unable to trigger blood-feeding behavior or activate all blood-sensitive stylet neurons. If activation of multiple stylet neuron subsets is required to initiate blood-feeding, it should decrease the possibility that a female accidentally engorges on nectar instead of blood. For instance, 298 mM glucose occasionally activated blood-sensitive neurons, but females still rejected this meal in the blood-feeding assay. How is information from this network of blood-sensitive neurons integrated to form the perception of blood? Experiments in *Drosophila melanogaster* and mice found that information from each taste quality remains segregated as each population projects to discrete regions in central taste-processing centers and activates different higher order neuronal populations (Chen et al., 2011; Harris et al., 2015; Lee et al., 2017; Marella et al., 2006). It will be interesting to see how signals from blood-sensitive subpopulations converge and if this processing stream is segregated from other sub-populations of stylet sensory neurons that do not respond to blood.

Prior to tasting blood, females must seek out a host. A previous study showed integration across sensory modalities like olfaction and thermosensation is critical to attract females to a blood source (McMeniman et al., 2014). Our biteOscope experiments further clarify that co-presentation of heat and CO_2_ specifically results in piercing, but not engorgement (Hol et al., 2020). Taken together, these results demonstrate that *Ae. aegypti* employ multimodal integration across various scales: within individual neurons, across taste qualities, and across sensory modalities. We speculate that integration increases specificity in the complex task of locating a suitable blood meal.

### Downstream circuits to control meal size and destination

Once ingestion begins, which mechanisms determine the meal size and destination associated with each feeding program? In *Drosophila melanogaster*, sensory information is relayed to various higher order neuronal populations inside and outside of the subesophageal zone (Scott, 2018). The neuronal populations identified thus far are taste quality-specific and are thought to ultimately communicate with subesophageal zone motor neuron populations that control pumping and ingestion (McKellar, 2016; Scott, 2018; Yapici et al., 2016). We show that sensory neurons from the stylet and labium project to distinct subesophageal zone regions. Does sensory input from the stylet and labium remain segregated as specialized blood-feeding and nectar-feeding circuits? Detection of blood by the stylet could be hardwired to dedicated sensory processing pathways, motor neurons, and muscles that control blood meal size and destination. Alternatively, information about blood and nectar could eventually converge onto overlapping neurons and activate motor neurons that generically initiate pumping. In this case, additional downstream input would be required to specify the appropriate meal size and digestive organ destination. The ability to implement circuit tracing techniques (Fosque et al., 2015; Matthews et al., 2019; Ruta et al., 2010; Talay et al., 2017) in *Ae. aegypti* will enable future studies of downstream central and peripheral circuits.

### The stylet is specialized to detect blood over nectar

The needle-like anatomy of the stylet is ideally adapted to blood-feeding (Choumet et al., 2012; Ramasubramanian et al., 2008) and we discovered that its functional properties directly encode a distinction between blood and nectar. We propose that specialization of peripheral sensory neurons in the stylet may explain why sugars do not promote nectar-feeding in the context of blood-feeding. This mechanism is distinct from previously described examples of food source valence changes upon nutrient deprivation or mating in *Drosophila melanogaster*. These cases normally involve a state-change that modulates the sensitivity of sensory neurons, and/or their downstream processing, to a given ligand (Devineni et al., 2019; Inagaki et al., 2012; Steck et al., 2018; Walker et al., 2015). One key difference between *Drosophila melanogaster* and *Ae. aegypti* feeding is that *Ae. aegypti* have two distinct feeding appendages. In *Ae. aegypti*, the stylet, but not the labium, contacts blood during blood-feeding and our *Gr4*>*TRPV1* chemogenetic experiments demonstrate that stylet neuron activity is dispensable for nectar-feeding. Therefore, we speculate that feeding appendage segregation and specialization is a mechanism to ensure the female ingests blood and not nectar in the context of blood-feeding. Furthermore, female-specific stylet sensilla are conserved across blood-feeding mosquito species and are absent in non-blood-feeding *Toxorhynchites* species (Lee and Craig, 1983). Although mosquito species differ in the minimum blood components required to initiate blood-feeding (Galun, 1987), blood detection via stylet neurons may be a conserved mechanism across many blood-feeding mosquito species. Understanding blood detection is fundamental to intervene in blood-feeding behavior, which is responsible for vector-borne disease transmission to millions worldwide.

## Supporting information

Jove Video 1

Jove Video 2

Jove Video 3

Jove Video 4

Jove Video 5

## SUPPLEMENTAL INFORMATION

Supplementary Figures S1-S5 and Videos 1-5 accompany the paper. All raw data and custom scripts in the paper are available at https://github.com/VosshallLab/Jove_Vosshall_2020.

## ACKNOWLEDGMENTS

We thank Josie Clowney, Matthew DeGennaro, Emily Dennis, Marcelo Dietrich, Laura Duvall, Annie Handler, Richard Jové, Kevin Lee, Molly Liu, Benjamin Matthews, Paul Muller, Nilay Yapici, and members of the Vosshall Lab for comments on the manuscript; Takeshi Morita for providing guidance and technical assistance in the development of stylet GCaMP calcium imaging; Benjamin Matthews for discussions and advice on RNA-seq and transgenic strain design; Laura Duvall for advice on the Glytube assays; Gloria Gordon and Libby Mejia for expert mosquito rearing; and Anjali Pandey for help with transgenic mosquito rearing. We thank Winrich Freiwald, Paul Muller, and Vanessa Ruta for discussions; Daniel Gross and Jim Petrillo of the Rockefeller Precision Instrumentation Technology resource center for consultation and advice on calcium imaging hardware; Gavin Jeffries of Fluicell for optimizing the BioPen to deliver blood; Connie Zhao and Bin Zhang of the Rockefeller University Genomic Resource Center for help with RNA-seq library preparation; Alison North and Christina Pyrgaki of the Rockefeller University Bio-Imaging Resource Center for help with confocal and structured illumination image acquisition; Rob A Harrell II at the Insect Transgenesis Facility at the University of Maryland for CRISPR-Cas9 and transgene injections; and Tasos Siskoglou and Andrew Gerson of Morrell for their help interfacing calcium imaging acquisition with ligand delivery. We thank Benjamin Matthews, Meg Younger and the *Aedes* Toolkit Group for providing early access to unpublished strains, and David Tian for his assistance in generating the *Brp-QF2w* strain.

This work was supported in part by grant # UL1 TR000043 from the National Center for Advancing Translational Sciences (NCATS, National Institutes of Health (NIH) Clinical and Translational Science Award (CTSA) program. VJ was supported in part by NIH T32-MH095246. This work was supported in part by a grant to The Rockefeller University from the Howard Hughes Medical Institute through the James H. Gilliam Fellowships for Advanced Study program to VJ. This material is based upon work supported by the National Science Foundation Graduate Research Fellowship Program under Grant No. NSF DGE-1325261 to VJ. Any opinions, findings, and conclusions or recommendations expressed in this material are those of the author(s) and do not necessarily reflect the views of the National Science Foundation. The work was supported in part by a Career Award at the Scientific interface (CASI) from the Burroughs Wellcome Fund (FJJH), the European Union’s Horizon 2020 research and innovation programme under the Marie Skłodowska-Curie grant agreement No 841893 — PiQiMosqBite (FJJH), Jane Coffin Childs Postdoctoral Fellowship (TRS), Kavli Neural Systems Institute postdoctoral fellowship (TRS), NIDCD R00-DC012069 (CSM). Development of biteOscope and associated work was supported by NIH DP2-AI124336 New Innovator Award (MP) and USAID Grand Challenges: Zika and Future Threats Award (MP). MP is an HHMI-Gates Faculty Scholar of the Howard Hughes Medical Institute. CSM is a New York Stem Cell Foundation – Robertson Investigator. LBV is an investigator of the Howard Hughes Medical Institute.

## AUTHOR CONTRIBUTIONS

VJ carried out all experiments in the paper with the following exceptions: FJHH carried out experiments in Figure 2 and Video 1; ZG and VJ together created the *Ir7a-QF2, Gr4-QF2*, QUAS-*TRPV1* strains; ZG and VJ together carried out experiments in Figure 1I, Figure S1A-C, Figure 3A,B, Figure S2B,C,E,F, Figure 6A,B, and Figure 7E; ZG performed experiments in Figure 1E,F,J, Figure S1D, and Figure 7B,C; ZZ created the *brp-QF2w* strain with the assistance of David Tian; CSM supervised and collaborated with ZZ on the *brp-QF2w* strain; MP supervised and collaborated with FJHH on the biteOscope; TRS carried out experiments in Figure 6C-F and Figure S5A,B; TSC and VJ designed the analysis for live imaging data; TSC wrote the analysis pipeline code, and VJ performed data analysis; VJ and LBV together conceived the study, designed the figures, and wrote the paper with input from all authors.

## DECLARATION OF INTERESTS

The authors declare no competing interests.

## MATERIALS AND METHODS

### Human and animal ethics statement

Blood-feeding procedures with live mouse and human hosts were approved and monitored by The Rockefeller University Institutional Animal Care and Use Committee (IACUC protocol 17108) and Institutional Review Board (IRB protocol LV-0652), respectively. Human subjects gave their written informed consent to participate.

### Mosquito rearing and maintenance

*Ae. aegypti* wild-type and genetically-modified strains were maintained and reared at 25 - 28°C, 70–80% relative humidity with a photoperiod of 14 hours light: 10 hours dark (lights on at 7 a.m.) as previously described (DeGennaro et al., 2013). Adult mosquitoes were provided constant access to 10% sucrose. Adult females were blood-fed on mice for stock maintenance, and occasionally on human subjects in the early stages of generating genetically modified strains. All mosquitoes were mated prior to experiments. Female mosquitoes were fasted for 14 - 24 hours in the presence of a water source prior to behavioral experiments.

### Generation of genetically modified mosquito strains

All new strains generated in this paper, except for *Brp-QF2w*, were injected into wild-type Liverpool embryos. *Brp-QF2w* was injected into wild-type Orlando embryos. All CRISPR-Cas9 and transgene injections were carried out at the Insect Transformation Facility at the University of Maryland Institute for Bioscience & Biotechnology Research. For instances where a transgene was integrated into the genome using homologous recombination, proper payload integration was confirmed in a strain using polymerase chain reaction (PCR). Animals were then back-crossed to wild-type Liverpool for at least three generations before crossing to corresponding QF2 or QUAS for experimental use. Details of plasmid construction are below. All homology arms for homology-directed integration were isolated by PCR from genomic DNA isolated from the Liverpool strain, except for *Brp-QF2w*, which was derived from the Orlando strain. When Gibson assembly was utilized in plasmid construction, oligonucleotide sequences are displayed in lower case to indicate homology to the adjacent fragment and upper case to indicate the target sequence.

#### 3xP3-eYFP-SV40-15xQUAS-dTomato-T2A-TRPV1-SV40

This plasmid was generated using NEBuilder HiFi DNA Assembly (New England Biolabs #E5520S), using the following fragments generated by PCR from the indicated template with the indicated primers:

1. Plasmid backbone with pBAC arms from *15xQUAS-dTomato-T2A-GCaMP6s* (Addgene plasmid #130666) (Matthews et al., 2019) (Primers: Forward, 5’-GATCTTTGTGAAGGAAC-CTTACTTCTGTGGTGTG-3’; Reverse, 5’-ATCCCCCGGGCTG-CAGGA-3’)
2. *QUAS-dTomato-T2A* from *15xQUAS-dTomato-T2A-GCaMP6s* (Primers: Forward, 5’-tcaatgtatcttaACTAGAGCGGCCGCCACC-3’; Reverse, 5’-cccgttgttccatAGGGCCGGGATTCTCCTC-3’)
3. *3xP3-eYFP-SV40* with YFP open reading frame from Addgene plasmid #62291 (Primers: Forward, 5’-atcgaattcctgcagcccgggggatGTTCCCACAATGGTTAATTC-3’; Reverse, 5’-ggccgctctagtTAAGATACATTGATGAGTTTGG-3’).
4. *Rattus norvegicus* TRPV1 (Genbank accession NM_031982.1) from *ASH:TRPV1* (Bargmann Lab plasmid #10.33.42, with permission from Dr. David Julius of UCSF) (Tobin et al., 2002) (Primers: Forward, 5’-aatcccggccctATGGAACAACGGGCTAGC-3’; Reverse, 5’-gaagtaaggttccttcacaaagatcACCCAGATAAC-GTCAACC-3’).

200 embryos were injected with 200 ng/µL plasmid and 200 ng/µL pBAC mRNA. Two independent transgenic lines were recovered, one of which was sex-linked. In pilot experiments, both lines showed qualitatively similar behavioral effects in the *Gr4*>*TRPV1* capsaicin experiments. All subsequent behavior and expression pattern experiments were performed using the non-sex-linked line.

#### Gr4, Ir7a, and Ir7f QF2 strains

These knock-in/knock-out strains were generated through CRISPR-mediated homologous recombination of the QF2 transcription factor (Potter et al., 2010; Riabinina et al., 2015) into the endogenous locus of the *Ae. aegypti* genome. *In vitro* transcription was performed using HiScribe Quick T7 kit (New England Biolabs #E2050S) following the manufacturer’s directions and incubating for 3 hr at 37°C. Following transcription and DNAse treatment for 15 min at 37°C, sgRNA was purified using RNAse-free SPRI beads (Ampure RNAclean, Beckman-Coulter #A63987), and eluted in Ultrapure water (Invitrogen #10977–015). For each line, 2000 embryos were injected with 300 ng/µL plasmid, 600 ng/µL Cas9 protein, and 40 ng/µL sgRNA. sgRNA DNA template was prepared by annealing oligonucleotides as previously described (Kistler et al., 2015). For all plasmids, fragments were generated by PCR from the indicated template with the indicated primers and assembled using NEBuilder HiFi DNA Assembly as detailed below.

#### Gr4-T2A-QF2 -SV40-3xP3-dsRed

1. Plasmid backbone from pUC19 (Primers: Forward, 5’-CTA-GAGTCGACCTGCAGGC -3’; Reverse, 5’-CCCGGGTAC-CGAGCTCGA -3’).
2. *Gr4* left homology arm (NCBI LOC5563657) (Primers: Forward, 5’-agtgaattcgagctcggtacccgg-gACTCTCCTAAAATCTCAAGTATAC-3’; Reverse, 5’-tctgccctctccTGCACGTTTGGGATACTTG-3’).
3. *Gr4* right homology arm (NCBI LOC5563657) (Primers: Forward, 5’-caatgtatcttaCAGGGAAAACTGGATCCATG-3’; Reverse, 5’-ttgcatgcctgcaggtcgactctagGTGTATTTGGAGCCTCAG-3’).
4. *T2A-QF2-SV40-3xP3-dsRed* with QF2 and dsRed open reading frame from *ppk301-T2A-QF2* (Addgene plasmid #130667) (Matthews et al., 2019) (Primers: Forward, 5’-tcccaaacgtgcaG-GAGAGGGCAGAGGAAGTC-3’; Reverse, 5’-ccagttttccctgTAA-GATACATTGATGAGTTTGGACAAAC-3). The sgRNA targeted exon 2 of the *Gr4* locus, target sequence with protospacer adjacent motif (PAM) underlined: GTATCCCAAACGTGCAAC-CA<u>GGG</u>.

#### Ir7a-T2A-QF2 -SV40-3xP3-dsRed

1. Plasmid backbone from pUC19 (Primers: Forward, 5’-cgatcaactataaCTAGAGTCGACCTGCAGGC -3’; Reverse, 5’-aatttgctttttaCCCGGGTACCGAGCTCGA-3’.
2. *Ir7a* left homology arm (Primers: Forward, 5’-cggtacccggg-TAAAAAGCAAATTTCACCATG-3’; Reverse, 5’-tctgccctctccATATACGTGACCCCAAATATC-3’).
3. *Ir7a* right homology arm (Primers: Forward, 5’-caatgtatcttaATCCAGAACGGGTGCGGTAG-3’; Reverse, 5’-ggtcgactctagTTATAGTTGATCGAGGAATTTCCGAATCC-3).
4. T A-QF2-SV40-3xP3-dsRed with QF2 and dsRed open reading frame from *ppk301-T2A-QF2* (Addgene plasmid #130667) (Matthews et al., 2019) (Primers: Forward, 5’-gggtcacgtatatGGA-GAGGGCAGAGGAAGTC-3’; Reverse, 5’-acccgttctggatTAA-GATACATTGATGAGTTTGGACAAAC-3’). The sgRNA targeted exon 1 of the *Ir7a* locus, target sequence with PAM underlined: TGGGGTCACGTATATCCAAA<u>TGG</u>.

*Ir7a* was not annotated in the AaegL5 NCBI RefSeq Annotation version 101 (Matthews et al., 2018). Genomic coordinates (NC_035107.1:37734383-37736188) were identified using the manual chemoreceptor annotation (Matthews et al., 2018). See the “Expression data and differential expression analysis” section below for additional annotation information.

#### Ir7f-T2A-QF2 -SV40-3xP3-dsRed

1. Plasmid backbone from pUC19 (Primers: Forward, 5’-atttt-gaggcgggCTAGAGTCGACCTGCAGGC-3’; Reverse, 5’-aatcagccagtcaCCCGGGTACCGAGCTCGA-3’).
2. *Ir7f* left homology arm (NCBI LOC5565007) (Primers: Forward, 5’-ctcggtacccgggTGACTGGCTGATTAGCTCATCCTATATAAGAA-3’; Reverse, 5’-ctctgccctctccACGCTCGCCACGCATCGA-GAAACACCCGG-3’).
3. *Ir7f* right homology arm (NCBI LOC5565007) Primers: Forward, 5’-tcaatgtatcttaTGTCGGTGATGAGGTCCAG -3’; Reverse, 5’-aggtcgactctagCCCGCCTCAAAATGTGCAC-3’).
4. T2A-QF2-SV40-3xP3-dsRed with QF2 and dsRed open reading frame from ppk301-T2A-QF2 (Addgene plasmid #130667) (Matthews et al., 2019) (Primers: Forward, 5’-gcgtggcgagcgtGGA-GAGGGCAGAGGAAGTC-3’; Reverse, 5’-ctcatcaccgacaTAA-GATACATTGATGAGTTTGGACAAAC-3’). The sgRNA targeted exon 1 of the *Ir7f* locus, target sequence with PAM underlined: GATGCGCGGTGAACGCATGT<u>CGG</u>.

#### Brp-QF2w strain

This knock-in strain was generated in an Orlando strain background using CRISPR-mediated homologous recombination of the QF2w transcription factor (Potter et al., 2010; Riabinina et al., 2015) into the endogenous *bruchpilot* locus (NCBI LOC5570381) of the *Ae. aegypti* genome. Briefly, a sgRNA (target sequence, with PAM underlined: GCAACTGGTACAGATGACAC<u>AGG</u>) targeting exon 21 of the *Brp* gene was generated using the HiScribe T7 Kit (New England Biolabs #E2040S) by incubating for 6 hours at 37°C. Following transcription and DNAse treatment for 15 min at 37°C, sgRNA was purified using RNAse-free SPRI beads (Agencourt RNAclean XP, Beckman-Coulter #A63987), and eluted in Ambion nuclease-free water (Life Technologies, #AM9937).The *Brp-T2A-QF2w-Hsp70-3XP3-dsRed* donor plasmid was constructed using the InFusion HD Kit (Clontech, #638910). Homology arms of ∼1 kb flanking the sgRNA cut site were cloned by PCR using template genomic DNA from the Orlando strain. To preserve the *Brp* coding sequence, the final 46 codons of *Brp* downstream of the cut site were included in the donor plasmid 5’ of the T2A motif. This 46 codon fragment was synthesized with synonymous codon substitutions to protect the sequence from Cas9 cleavage and to minimize homology between the plasmid insert and the targeted locus (IDT, gBlocks). An injection mix containing sgRNA (80 ng/µL), donor plasmid (700 ng/µL) and Cas9 protein (300 ng/µL, PNA Bio #CP01-200) was injected into 1533 Orlando strain embryos.

#### Brp-T2A-QF2w-Hsp70-3XP3-dsRed

1. Plasmid backbone from psL1180, linearized with restriction enzymes NsiI-HF (New England Biolabs #R3127S) and AvrII (New England Biolabs #R0174S).
2. *Brp* left homology arm (NCBI LOC5570381) (Primers: Forward, 5’-caggcggccgccataATGACCGGCTACCATGACCAC-TTTATAGTA -3’; Reverse, 5’-TCATCTGTACCAGTTGCAG-TAAACGTTCC -3’).
3. *Brp* right homology arm (NCBI LOC5570381) (Primers: Forward, 5’-tgtatcttatcctagCACAGGAAGAGCAGAACCAAAAA-GAAAAGAC -3’; Reverse, 5’-tattaataggcctag-TTTCGAATCTGTGACAAATTTCCCGATAAGAACT -3’).
4. QF2w-Hsp70 from pQF2wWB (Addgene plasmid #61313) (Primers: Forward, 5’-GCAAAACGCTTAACGCTGCG -3’, Reverse, 5’-cgtaggataacttcgGGATCTAAACGAGTTTTTAAGCAAACT -3’)
5. *Brp* synthetic fragment with synonymous codon substitutions: AACTGGTACAGATGACCCAGGAAGAACAGAAC-CAGAAGGAAAAGACCATCATGGATCTGCAG-CAGGCCCTGAAGAACGCCCAGGCCAAGCTGAAGACCGCCCAGTCGCAGCCGCAGGATGCCGGACCGGCCGGAT-TCCTGAAGTCGTTCTTTGGATCGGGAGAGGG

### Ligands for feeding experiments

#### Sheep blood

(Hemostat Laboratories #DSB100) was used within 1 week of arrival.

#### Nucleotides

ATP (Adenosine 5′-triphosphate disodium salt hydrate, Sigma #A6419), AMP-PNP (β,γ-imidoadenosine 5’-triphosphate lithium salt hydrate, Millipore Sigma #10102547001), AMP-CPP (α,β-methyleneadenosine 5’-triphosphate lithium salt, Jena Bioscience #NU-421-25), AMP-PCP (β,γ-Methyleneadenosine 5′-triphosphate disodium salt, Millipore Sigma #M7510). ATP and non-hydrolyzable analogues were reconstituted and aliquoted in 25 mM NaHCO_3_.

#### Sugars

sucrose (Fisher Scientific #S5-3), cellobiose [D-(+)-cellobiose, Millipore Sigma #22150], fructose [D-(-)-Fructose, Millipore Sigma #F0127], glucose [D-(+)-Glucose, Millipore Sigma #G7528].

#### Additional blood components

NaCl (Millipore Sigma #S6546), NaHCO_3_ (Fisher Scientific #S233), albumin (human serum, Millipore Sigma #A9511), hemoglobin (human, Millipore Sigma #G4386), gamma-globulin (human blood, Millipore Sigma, #H7379).

#### Capsaicin

(E)-capsaicin (Tocris #0462)

### Blood-feeding assay (Glytube)

7 to 21 day-old female mosquitoes were anesthetized at 4°C and sorted into groups of 15-20 females, and placed into a 32 oz. HDPE plastic cup (VWR #89009-668). The cup was prepared by cutting a 10 cm hole in the lid with a razor blade, covering the cup with a 20 cm x 20 cm piece of white 0.8 mm polyester mosquito netting (American Home & Habit Inc. #F03A-PONO-MOSQ-M008-ZS) and securing the mesh to the cup by snapping on the modified lid. Animals recovered overnight at 25 - 28°C, 70–80% relative humidity with access to water. The assay chamber was a modification of previously published methods (McMeniman et al., 2014) and used a translucent polypropylene storage box 36 cm L x 31 cm W x 32 cm H with a removable lid. One 1.5 cm hole made on the chamber wall allowed silicone tubing for CO_2_ delivery. The CO_2_ diffusion pad (8.9 cm x 12.7 cm; Tritech Research) was affixed to the inner center of the lid to allow delivery of purified air and CO_2_ to condition the chamber atmosphere during the trial. Up to 4 cups were placed in the chamber per trial and feeding positions were randomized according to meal during assays. Females were fed sheep blood or test ligands using Glytube membrane feeders exactly as described (Costa-da-Silva et al., 2013), except the Parafilm feeding surface was not rubbed on human skin prior to offering the Glytube to mosquitoes to avoid introducing contact chemosensory cues as secondary stimuli in our experiments. In Figure 1, Figure 2 and Figure S1, the saline meal contained 110 mM NaCl and 20 mM NaHCO_3_. All meals and Glytubes were preheated for at least 15 min in a 45°C water bath and, if required, ATP or non-hydrolyzable ATP analogues were added to meals immediately before feeding and mixed by vortexing. At the start of each trial, cups were placed in the assay chamber and allowed to acclimate for 5 min before 1 Glytube with 1.5 mL of each meal was placed on top each cup and CO_2_ was turned on for 15 min. In Figure 1M, Figure S1E,G,I, and Figure 5A-B, fed females were scored by eye for engorgement of the abdomen. In the rare cases that females were considered partially fed they were counted as non-fed and discarded. To sample the weights of these females (Figure 1K,L and Figure S1F,H,J), a selection of engorged individuals was weighed in groups of 5 females and the resulting weight in mg was divided by 5 to report the average weight per female. In Figure 1E and Figure 7C, Glytube feeding was performed as above, except that fluorescein (Amresco #0681) was added as a fluorescent tracer to each meal (blood, sucrose, fructose, glucose, or water) at a final concentration of 0.002%. After feeding, females were stored at -20°C until fluorescence reading. A 96-well PCR plate was prepared with one 3 mm diameter borosilicate solid-glass bead (Millipore Sigma #Z143928) and 100 µl PBS in each well. 8 wells were used to generate a reference standard curve. These wells contained a single unfed mosquito and the following volumes of the same fluorescent meal fed to test mosquitoes: 5, 2.5, 1.25, 0.625, 0.3125, 0.15625, 0.078125, or 0 µL. One test group mosquito was added to each of the remaining wells. Tissue was disrupted using TissueLyser II (Qiagen) and briefly centrifuged at 2000 rpm for 1 – 2 min. 20 µL of tissue lysate was added to 180 µL PBS in a black 96-well plate (ThermoFisher #12-566-09). Fluorescent intensity for each well was measured using the 485/520 excitation/emission channel of a Varioskan Lux (ThermoFisher #VL0000D0) plate reader. Using the reference dilution curve, fluorescent measurements were converted to volume (µL) of solution ingested. Measurements below the level of detection were quantified as 0 for plotting and statistical analysis.

### Nectar-feeding assay

Animals were prepared exactly as described for the Glytube assay above. Consumption of nectar was quantified by doping the meal with 0.002% fluorescein. A cotton ball (Fisher Scientific #22456880) was soaked in each test meal, dabbed on a Kimwipe to prevent excess liquid from dripping through the mesh, and placed on top of the mesh covering the cup. Animals were allowed to feed for 4 hours. After feeding, animals were frozen at -20°C and fluorescence reading was performed as described above.

### Meal size quantification

In Figure 1E, F, we analyzed the average meal size of mosquitoes that fed on blood or sugar respectively. We therefore excluded mosquitoes that did not feed from the meal size analysis. To set a cut-off for whether or not a mosquito fed, we included unfed control groups that were not offered a meal and should be a true 0. We detected fluctuations in baseline from 0 – 0.0304 µL. We therefore set a cut-off at 0.05 µL and excluded animals in the blood or sugar experimental group that measured < 0.05 µL. We then applied this 0.05 µL cut-off for statistical analysis in subsequent meal size quantification experiments in Figure 1J, Figure S1D, and Figure 7B,C: all values < 0.051 were replaced with 0.05.

### Chemogenetic capsaicin feeding assay

Chemogenetic experiments using capsaicin to activate *Gr4*>*TRPV1* sensory neurons were carried out exactly as the nectar-feeding experiments described above except that 50 µM capsaicin in 0.1% DMSO or 0.1% DMSO only-control was added to the meals.

### biteOscope assay

Stylet piercing behavior was characterized using the biteOscope (Hol et al., 2020). Briefly, all meals were prepared exactly as for the Glytube experiments above. The meal was applied on the rectangular section on the outside of a 70 mL Falcon cell culture flask and covered with parafilm. To maintain meal temperature, the flask was filled with warm water maintained at 37°C using a Raspberry Pi controlled Peltier element. The flask was mounted in the floor of a 10 cm x 10 cm x 10 cm acrylic cage. A camera (Basler #acA2040-90um) and two white LED arrays for illumination (Vidpro #LED-312) were mounted outside the cage to image mosquitoes interacting with the bite substrate. At least 12 hours prior to the experiment, females were given water instead of 10% sucrose. At the start of each trial, an individual female was introduced into the cage and the experimenter blew on the cage 2 times 10 sec to provide human cues. Images were acquired at 10 frames/sec using Basler Pylon 5 software running on Ubuntu 18.04. Each female was recorded for 700 sec regardless of engorgement status. Images were processed using custom code written in Python (available from Github: https://github.com/felixhol/biteOscope) using SciPy (Virtanen et al., 2019), TrackPy (Allan et al., 2019), and OpenCV (Bradski, 2000) packages to determine the presence and location of a mosquito. Engorgement status of a mosquito was determined by measuring abdominal size by fitting an active contour model to its abdomen. Stylet piercing events were scored by manual visual analysis of the images.

### TO-PRO-3 staining

7 to 14-day old animals were anesthetized on ice. Tissue fixation followed modification of previously published methods (Matthews et al., 2019) as follows. Heads were carefully removed from the body by pinching at the neck with sharp forceps. Heads were placed in a 1.5 mL tube for fixation with 4% paraformaldehyde (PFA), 0.1 M Millonig’s Phosphate Buffer (pH 7.4), 0.25% Triton X-100, and nutated for 3 hour at 4°C. Samples were dissected and samples of the same tissue were grouped into a cell strainer cap (Fisher Scientific #08-771-23) that was cut to fit into 1 well of a 24-well plate containing PBS with 0.25% Triton X-100 (PBT). All subsequent steps were performed on a low-speed orbital shaker at room temperature. Samples were washed at least 5 times 20 min and transferred to 0.25% PBT with 1:400 TO-PRO-3 (ThermoFisher #T3605) for 2 nights. Samples were washed at least 5 times 20 min in 0.25% PBT. After washing, tissues were briefly transferred to a well of SlowFade diamond (ThermoFisher #S36972) to eliminate excess PBT. Samples were then mounted in SlowFade. Within each experiment, all image acquisition parameters were maintained across both sexes.

### dTomato visualization

7 to 14 day-old mosquitoes were anesthetized on ice. Heads were fixed and samples were dissected and washed as described above. After washing, tissues were briefly transferred to a well of Vectashield (Vector Laboratories #H-1000) to remove excess PBT. Samples were then mounted in Vectashield. Within each genotype, all image acquisition parameters were maintained across tissue types. At higher laser power, we observed faint cells in *Ir7f*>*dTomato-T2A-GCaMP6s* female labiums (Figure 6B, right panel) but we suspect that they are not neurons because we did not observe nerve fibers exiting the labium or projecting to the posterior subesophageal zone where labium neurons normally terminate (Figure 3L, Figure S2H).

### Phalloidin, DAPI, and FITC staining

7-14-day old mosquitoes were anesthetized on ice. Stylets were dissected and placed directly into a 24 well-plate containing 4% paraformaldehyde, 0.1 M Millonig’s Phosphate Buffer (pH 7.4), 0.25% Triton X-100. All subsequent steps were performed on a low-speed orbital shaker at room temperature. Samples were washed at least 4 times 15 min before placed overnight in permeabilization solution from the previously published iDISCO method (Renier et al., 2014). Samples were then incubated in iDISCO PTwH solution (for 1L: 100 mL 10x PBS, 2mL Tween-20, 1 mL of 10mg/mL Heparin stock solution) with 5% DMSO for at least 2 nights at room temperature with the following reagents: (1) 1:20 AlexaFluor 594 phalloidin (ThermoFisher #A12381) (Figure 3F) or (2) 1:20 AlexaFluor 488 phalloidin (ThermoFisher #A12379) and 1:500 DAPI (Millipore Sigma #D9542) (Figure S2D) or (3) 1:20 AlexaFluor 647 phalloidin (ThermoFisher #A22287) (Figure S2F) or (4) 2 mg/mL FITC (Millipore Sigma #1.24546) (Figure S2A). Samples were then washed at least 4 times 15 min and mounted in Vectashield, except samples containing AlexaFluor 647, which were mounted in SlowFade.

### Dextran dye-fills

7 to 14 day-old mosquitoes were anesthetized on ice. The labium was separated from the stylet using forceps. Mosquitoes were affixed on their side to a plastic dish (Falcon #353001) using UV-curable glue (Bondic, Amazon #B0181BEHQU) or double-sided tape so that the stylet and labium were flat on the dish and distal tips were separated. For stylet dye-fills, a scalpel was used to cut approximately 300-750 µm away from the distal tip and 1 µL of Dextran, Texas Red, 3000 MW, Lysine Fixable (ThermoFisher #D3328) diluted to 1 mg/10 µL in External Saline was added immediately. The External Saline recipe (Matthews et al., 2019) is based on *Drosophila melanogaster* imaging saline: 103 mM NaCl, 3 mM KCl, 5 mM 2-[Tris(hydroxymethyl)methyl]-2-aminoethanesulfonic acid (TES), 1.5 mM CaCl_2_, 4 mM MgCl_2_, 26 mM NaHCO_3_, 1 mM NaH_2_PO_4_, 10 mM trehalose, 10 mM glucose, pH 7.3, osmolality adjusted to 275 mOsm/kg. The mosquito was left on ice and covered for approximately 3-5 min before excess dye was pipetted up. Mosquitoes were left at 4°C overnight with a moist Kim-wipe to prevent desiccation. Heads were then removed and fixed as described above. For double fills, the mosquito was prepared as described above. The labium was cut at the base of the labellar lobes using a scalpel and 1 µL of Dextran, Texas Red diluted to 1 mg/10 µL in External Saline was added immediately. The mosquito was left on ice and covered for approximately 3-5 min before excess dye was pipetted up. The stylet was cut approximately 300 – 750 µm away from the distal tip and 1 µL of Dextran, Fluorescein and Biotin, 3000 MW, Lysine Fixable (ThermoFisher #D7156) diluted to 1 mg/10 µL in External Saline was immediately added. The mosquito was left on ice and covered for approximately 3-5 min before excess dye was pipetted up. Mosquitoes were left at 4°C overnight with a moist Kimwipe to prevent desiccation. Heads were then removed and fixed as described above. Fixed heads were then dissected and brains were placed in cell-strainer caps (Falcon #352235) in a 24 well-plate. Brains were stained using a modification of previously published methods (Matthews et al., 2019). All subsequent steps were performed on a low-speed orbital shaker. Brains were washed at room temperature in PBT for at least 4 times 15 min. Brains were permeabilized with 4% Triton X-100 with 2% normal goat serum (Jackson ImmunoResearch #005-000-121) in PBS at 4°C for 2 days. Brains were washed at least 5 times 15 min with PBT at room temperature. Brains were incubated in PBT plus 2% normal goat serum with primary antibodies at the following dilutions: rabbit anti-fluorescein (ThermoFisher #A889) 1:500 and mouse anti-*Drosophila* Brp (nc82) 1:50. The nc82 hybridoma developed by Erich Buchner of Universitätsklinikum Würzburg was obtained from the Developmental Studies Hybridoma Bank, created by the NICHD of the NIH and maintained at The University of Iowa, Department of Biology, Iowa City, IA 52242. Primary antibodies were incubated for 3 nights at 4°C degrees then washed at least 5 times 15 min with PBT at room temperature. Brains were incubated with secondary antibody for 3 nights at 4°C with secondary antibodies at 1:500: goat anti-rabbit Alexa Fluor 488 (ThermoFisher #A-11008) and goat anti-mouse Alexa Fluor 647 (ThermoFisher #A-21236). Brains were then washed and mounted in Vectashield.

### Brain immunostaining

to 9-day old mosquitoes were anaesthetized on ice. Brains were fixed, dissected, washed and permeabilized as described above. Primary antibodies were used at the following dilutions: rat anti-mCD8 (Invitrogen #14008185) 1:100, and a concentrated aliquot of mouse anti-*Drosophila* Brp 1:5000. Brains were then washed 5x for at least 30 min at room temperature. Brains were then incubated with secondary antibodies in PBT with 2% normal goat serum for 2 days at 4°C. The following secondary antibodies were used at 1:500 dilutions: goat anti-rat Alexa Fluor 647 (Invitrogen #A21247) and goat anti-mouse Alexa Fluor 555 (Invitrogen #A32727). Brains were then washed 6 times in PBT at room temperature for at least 30 min then mounted in SlowFade diamond. Within each genotype, all image acquisition parameters were maintained across both sexes. 3xP3 was used as a promoter to mark transgene insertion as previously described (Matthews et al., 2019). To avoid any interference from possible 3xP3 signal, we used a different laser excitation/secondary antibody for monitoring *Ir7a, Ir7f*, and *Gr4* expression.

### Confocal image acquisition

Images were acquired with a Zeiss Axio Observer Z1 Inverted LSM 880 NLO laser scanning confocal microscope (Zeiss) with a 25x/0.8 NA immersion-corrected objective at a resolution of 2048 x 2048 or 1024 x 1024 pixels. When necessary, tiled images were stitched with 10% overlap. Confocal images were processed in ImageJ (NIH).

### Ex-vivo stylet prep for calcium imaging

Calcium imaging was performed on an inverted Ti-2E wide-field microscope (Nikon) with a dual FITC/TRITC bandpass cube and alternating emission wheel with 520/40 GFP and 628/40 RFP bandpass filters. A nd2 filter was added with the 628/40 RFP bandpass filter to attenuate dTomato signal. Images were acquired with a 25x/0.9 N.A. water-immersion objective (Nikon) and Zyla 4.2 Plus camera. Calcium imaging experiments were performed on female mosquitoes that were 7–14 days post-eclosion. Prior to dissection, the imaging chamber was prepared by affixing a Gold Seal Cover Glass, No. 1 22 x 40 mm coverslip (Ted Pella #260353) to a recording chamber using silicone lubricant (Dow Molykote 111 O-Ring Silicone Lubricant). A fast exchange recording chamber (Warner Instruments #64-0230) was used for perfusion-only experiments and a low-profile large bath recording chamber (Warner Instruments #640236) was used to accommodate the BioPen apparatus. Silicone lubricant approximately 100-200 µm in diameter was placed slightly off-center on the coverslip. After preparing the chamber, females were anesthetized briefly at 4°C for dissection. The labium was removed to expose the stylet, and then the stylet was detached at the proximal end using a scalpel (Feather disposable scalpel, No. 11, Fisher Scientific #FH/CX7281A). The severed end was immediately placed in the silicone lubricant with the stylet tip facing the center of the coverslip. Great care was taken to place the stylet flat along the coverslip so that all stylet neurons could be imaged in one plane. This process often involved carefully removing the maxillae and mandibles without damaging the stylet. However, if the stylet was already flat, it was not necessary to remove additional appendages as they did not interfere with image acquisition. The most distal 300 µm of the stylet tip remained free of silicone lubricant to prevent interference with ligand delivery. Once the stylet was secured to the coverslip, the chamber was filled with MilliQ water and the perfusion and/or BioPen fluidics were inserted into the chamber. dTomato fluorescence was examined before and throughout imaging to verify that the stylet nerves were intact. The sample remained stable during the duration of the imaging session in all animals that were included in this study. Each image acquisition captured one GCaMP image and one dTomato image separated by less than the 100 ms required to switch the filter wheel. Image acquisition was triggered at a rate of approximately 2 frames per sec for each channel (2 sets of GCaMP/dTomato images per sec).

### Perfusion ligand delivery

Two independent ValveBank8 Pinch Valve perfusion systems (Automate Scientific #13-pp-54) with BubbleStop8 60 mL Syringe Heater (Automate Scientific #10-8-60-G) were automatically controlled by NIS-Elements software (Nikon). To ensure full perfusion chamber exchange, ligands were perfused for 30 sec followed by a 45 sec recovery period before the next ligand. Ligand delivery switched from water (baseline) to ligand of interest with the following exceptions. Since ATP is rapidly hydrolyzed in water, ATP was always delivered in a buffer of 25 mM NaHCO_3_. 25 mM NaHCO_3_ was delivered for 30 sec to establish a baseline, after which ATP dissolved in 25 mM NaHCO_3_ was applied. Responses above the baseline were considered ATP responses. In control experiments, we demonstrated that ATP dissolved in PBS activated these same neurons after pre-equilibration in PBS (see raw data at GitHub: https://github.com/VosshallLab/Jove_Vosshall_2020). In Figure 7F,G stylets were pre-equilibrated in 298 mM cellobiose for 30 sec prior to the isomolar sugar of interest to control for osmotic effects. 298 mM cellobiose was behaviorally inactive in both the blood- and nectar-feeding assays (Matthews et al., 2019) (see: https://github.com/VosshallLab/Jove_Vosshall_2020).

### Microfluidic ligand delivery using the BioPen

The BioPen tip holder (Fluicell) was secured using a MP-285 micromanipulator (Sutter #SU-MP-285). Each BioPen tip was prepared according to the manufacturer’s instructions with the following exceptions. First, the initial “New Tip” protocol was run with MilliQ water in each well to prime the microfluidic channels. Once the protocol was completed, water was removed from each BioPen well and replaced with test ligands. 0.0002% fluorescein was added to each test ligand to visualize the size and location of ligand delivery in each trial. For solutions containing NaHCO_3_, the fluorescein signal was much brighter, so 0.00002% fluorescein was used instead. For each ligand, the BioPen stimulus was ON for 20 sec with a 60 sec recovery before the next stimulus.

### Analysis of GCaMP6s data

All calcium imaging data were processed with Nikon Elements software. Regions of interest (ROIs) were selected based on the dTomato fluorescence intensity and used for analysis of GCaMP6s signal. Great care was taken to draw ROIs on the cell body of interest and not on *en passant* processes or slightly overlapping cell bodies. To exclude background noise, a cut-off of 0.25 peak ΔF/F_0_ was set as the minimum threshold for activation. This cut-off intentionally filters for clear activation and does not distinguish between background noise and weak activation. Occasionally (less than 1 cell body per animal) it was difficult to avoid the halo, especially if baseline GCaMP fluorescence was very low in a given cell body. In these rare cases, the cell body was not considered to be activated. All traces with sample motion, as determined by dTomato fluorescence instability, were discarded.

Once raw fluorescence values were extracted for each neuron/stimulus (ligand) pair, ΔF/F_0_ calculations were performed using a custom R script (R version 3.6.0) where ΔF/F_0_ = (F – F_0_)/F_0_. To determine the baseline fluorescence (F_0_) 5 frames (∼2 fps) were averaged before stimulus presentation. To determine peak F to a given stimulus, the average of 3 frames at the peak during stimulus delivery was determined for each stimulus. This process was repeated twice for each stimulus so that the peak ΔF/F_0_ value represented in all plots is the average peak ΔF/F_0_ for 3 independent stimulus presentations. Stimulus trains were delivered so that each stimulus was only presented once per trial. Therefore, the final value represents the average peak stimulus response collected from three trials. Once all averages had been calculated, the dataset from individual females were analyzed and represented in multiple ways. Heat maps in Figure 4G, Figure 5D, Figure 6G,H, and Figure S3A were generated using a custom R script. Each box represents average peak ΔF/F_0_ to a given stimulus as described above. The heat map color scale is log2 to increase dynamic range and the minimum and maximum color value was set to 0.25 and 3 respectively. Plots in Figure 4J, Figure 5E, and Figure 7G were plotted using Prism 8 (GraphPad) and a neuron was considered activated if peak ΔF/F_0_ > 0.25. Peak ΔF/F_0_ scatter plots in Figure 4H, Figure 5F, Figure 6J,L, and Figure 7H-J were generated using Prism 8 (GraphPad). Custom R script available at Github: https://github.com/VosshallLab/Jove_Vosshall_2020.

### Hierarchical clustering

In Figure 5G-I individual neuron data from the 5 individuals in Figure S3A were pooled and clustered by hierarchical clustering using Euclidean distance with complete linkage and visualized with the Pheatmap R package. Five clusters were derived as the optimal number of clusters from evaluating the percent of explained variance between and within putative clusters. Scatter plots (Figure 5H) and Box plots (Figure 5I) of peak ΔF/F_0_ for neurons within a given cluster were plotted in base R. In Figure 5I the statistical significance of responses across ligands was evaluated using the one-sample Wilcoxon signed rank test in the wilcox.test R package. A cluster was considered activated by a given ligand if p < 0.05 when compared to the hypothetical value 0.25: wilcox.test(GroupA_Values, mu = 0.25, alternative = “greater”). Custom R script available at Github: https://github.com/VosshallLab/Jove_Vosshall_2020.

### Tissue dissection and RNA extraction

7 to 11-day old mosquitoes were cold-anesthetized and kept on ice for up to 30 min or until dissections were complete. For labium samples, the labium was removed by forceps and immediately flash-frozen in DNA Lo-bind nuclease-free tubes (Fisher Scientific #13-698-790) contained in a CoolRack (Biocision #BCS0137) in dry ice for snap-freezing tissue. For female and male stylet samples, the labium was removed first. The stylet was detached half-way from the tip using a scalpel and immediately flash-frozen as described above. Extreme caution was taken during the tissue dissection and RNA extraction process to ensure that there was no contamination from other mosquito tissues or RNases. Each dish, forcep, and scalpel was carefully cleaned with 70% ethanol and RNase-away (ThermoFisher #7003) after every dissection or dissection attempt. Once the labium was removed, the stylet was discarded if there was any contact between the stylet and any surface other than the cleaned dish, forceps, or scalpel. A dedicated pair of stylet-only forceps was used to place the detached stylet into the collection tube. The following number of mosquitoes was used for each female library: female stylet, 25; male stylet, 25; female labium, 4. Each sample group was dissected in parallel to avoid artifacts and batch effects. Dissected tissue was stored at -80°C until RNA extraction. RNA extraction was performed using the PicoPure Kit (ThermoFisher #KIT0204) with the following exception for homogenizing tissue: instead of lysis buffer, 240 µL of TRIzol (ThermoFisher #15596018) was added to the collection tube on ice. Custom-order molecular biology grade, low-binding zirconium beads in 100 µm, 200 µm and 800 µm were used to disrupt tissue (OPS diagnostics). An RNase free spatula (Corning #CLS3013) was used to add 1 scoop each of 100 µm and 200 µm beads and ∼100 µL of 800 µm beads to collection tube. Tubes were briefly spun down in a tabletop centrifuge before disruption in a Tissue-Lyser II (Qiagen #85300) for 2 min 30 sec at 30 Hz. Tubes were briefly spun down again in tabletop centrifuge and returned to the TissueLyser II for an additional 2 min at 30 Hz. The remaining TRIzol extraction steps were performed in the hood according to manufacturer’s instructions: tubes stood at room temperature for 5 min before 48 µL of chloroform:isoamyl alcohol 24:1 was added (Sigma #C0549). Tubes were hand-shaken for 30 sec and left to stand for 2 min before centrifuging at 12,000 x *g* for 15 min at 4°C. The aqueous Trizol layer was then removed and added into the PicoPure column, up to 180 µL at one time. Subsequent steps were performed according to PicoPure manufacturer’s instructions, including DNase treatment.

### RNA-seq library preparation and sequencing

Labium samples were run on Bioanalyzer RNA Pico Chip (Agilent #5067-1513) to determine RNA quantity and quality and were used as a proxy for overall sample integrity because female and male stylet samples fell below the level of detection. Labium samples were diluted 1:10 before cDNA amplification to more closely approximate stylet samples. cDNA synthesis was performed using SMART-Seq v4 Ultra Low Input RNA Kit for Sequencing (Takara #634894) according to the manufacturer’s instructions except that 10 µL instead of 9 µL was used to optimize for low RNA input. The number of PCR amplification cycles was adjusted for each sample group based on the number of cycles needed to detect RNA in the lowest input sample as determined by Bioanalyzer High Sensitivity DNA Kit (Agilent #5067-4627). Negative controls for each group were run in parallel to ensure that additional cycles did not result in unspecific background product. All samples within one group were subjected to the same number of PCR amplification cycles. The female labium and female stylet samples underwent 20 and male stylet 22 cycles. The full-length cDNA output was processed with Nextera XT DNA library preparation kit (Illumina #FC-131-1024) according to manufacturer’s instructions. Library quantity and quality were evaluated using High Sensitivity DNA ScreenTape Analysis (Agilent #5067-5585) prior to pooling. Bar-coded samples from all tissues were pooled in an equal ratio before distributing the pool across 3 sequencing lanes. Sequencing was performed at The Rockefeller University Genomics Resource Center on a NextSeq 500 sequencer (Illumina). All reads were 1 x 75 bp. Data were demultiplexed and delivered as fastq files for each library. Sequencing reads have been deposited at the NCBI Sequence Read Archive (SRA) under BioProject PRJNA605870.

### Expression data and differential expression analysis

All reads were trimmed using TrimGalore version 0.4.2. (https://github.com/FelixKrueger/TrimGalore) with minimum read length of 35 base pairs. Reads from individual libraries were mapped to the AaegL5 genome (Matthews et al., 2018) using STAR version 2.5.2a (Dobin et al., 2013). A custom gene annotation was generated by merging AaegL5 with the more recent manual chemoreceptor annotation for *OR*s, *GR*s and *IR*s (Matthews et al., 2018). This merged annotation and the R script used to generate it is available at Github: https://github.com/VosshallLab/Jove_Vosshall_2020. For each of these chemoreceptors, the manual annotation replaced the AaegL5 RefSeq annotation. If the chemoreceptor did not previously exist in AaegL5 RefSeq, it was added. Reads mapping to each were mapped to transcript coding regions (UTRs and multi-mappers were excluded) using featureCounts version 1.5.0-p3 (Liao et al., 2014). For abundance visualization, raw counts were converted to TPM (see TPM table https://github.com/Voss-hallLab/Jove_Vosshall_2020). RNA-seq TPM plots were generated using ggplot2 version 3.2.0 (R Development Core Team, 2017) in RStudio R 3.6.0. Raw counts were used for differential expression analysis in R using DESeq2 version 1.24.0 (Love et al., 2014). Sweet GRs analyzed in Figure 7D were derived from a recent genome reannotation (Matthews et al., 2018). A previous study reported odorant receptor (*OR*) expression in the stylet (Jung et al., 2015). We found no evidence for this in our RNA-seq dataset, although we did detect *Orco* and *OR* expression in the labium (see https://github.com/VosshallLab/Jove_Vosshall_2020).

### Filtering for stylet-specific genes

To obtain the 53 genes enriched in the female stylet compared to the female labium and male stylet (Figure S4A,B), we examined TPM values for non-mouthpart tissues that were previously profiled in a comprehensive dataset (Matthews et al., 2018; Matthews et al., 2016). A transcript was considered female stylet-specific if the average TPM expression across a given tissue was < 0.5 TPM for all tissues profiled by Matthews and colleagues, except for the Probosics and Rostrum samples because these samples included mouthparts. To calculate average TPM, we used the most recent dataset aligned to the L5 genome and quantified using NCBI Ref-Seq Annotation version 101 (Matthews et al., 2018). If a transcript was present in the NCBI RefSeq annotation and the manual chemoreceptor annotation published alongside (Matthews et al., 2018), we used the TPM value quantified using the manual chemoreceptor annotation because the NCBI RefSeq annotation is missing a handful of chemoreceptors, including *Ir7a*.

### Quantification and statistical analysis

All statistical analysis was performed using Graphpad Prism Version 8 and RStudio R 3.6.0. Data collected as raw values are shown as mean±SEM or mean±SD. Details of statistical methods are reported in the figure legends.

### Data and software availability

All data in the paper with the exception of raw GCaMP data files are available on Github at https://github.com/Voss-hallLab/Jove_Vosshall_2020. All plasmids described in this paper are available at Addgene. Scripts for merged genome annotation and calcium imaging analysis are available on Github at https://github.com/VosshallLab/Jove_Vosshall_2020

## SUPPLEMENTARY FIGURES

**Figure S1:**
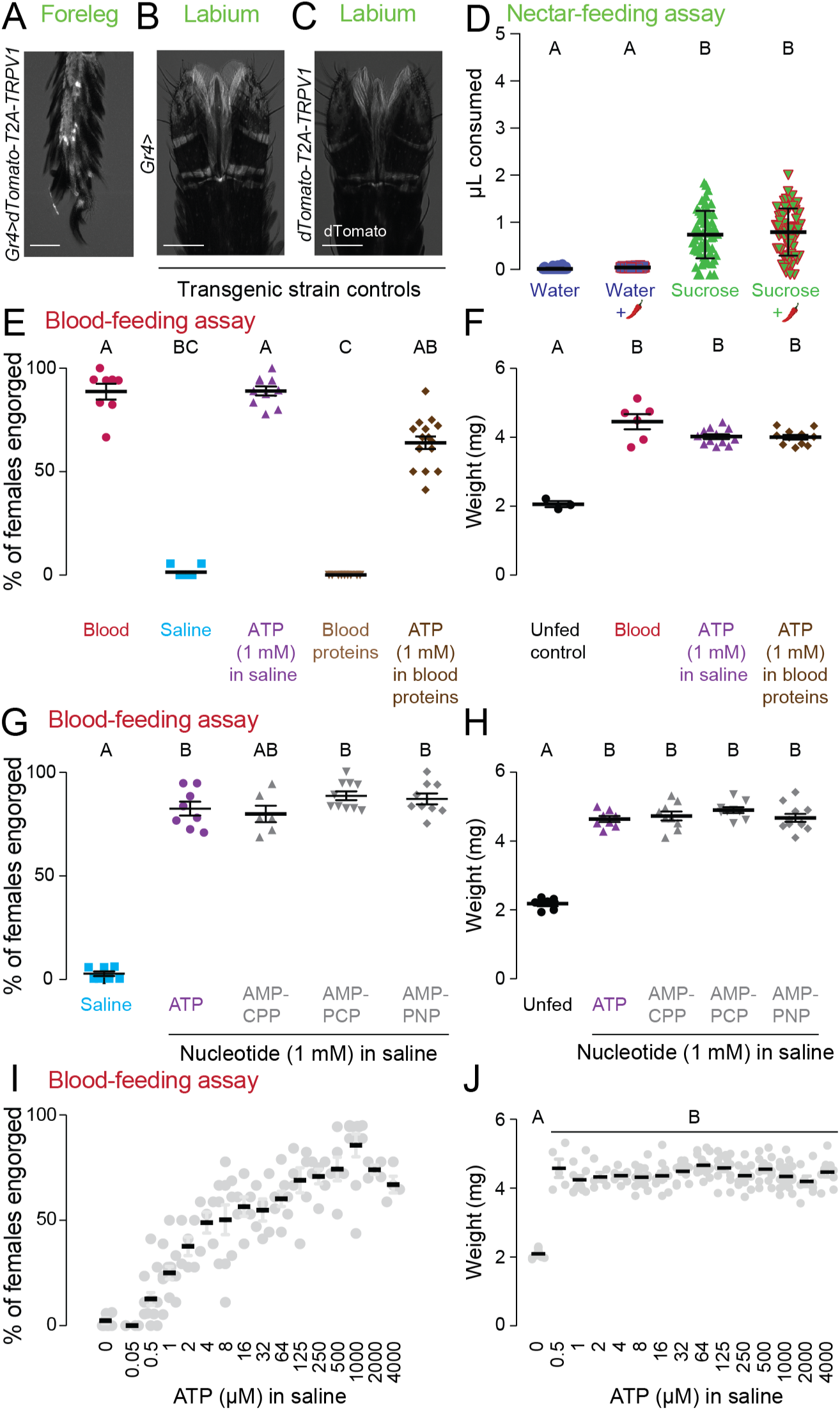
Features of blood- and nectar-feeding behavior (Related to Figure 1) (A-C) Confocal image of dTomato expression with transmitted light overlay in *Gr4*>*dTomato-T2A-TRPV1* foreleg (A), *Gr4* control labium (B), and *dTomato-T2A-TRPV1* control labium (C). Scale bar: 50 µm. (D) Volume of meal consumed by wild-type mosquitoes. Chili pepper indicates addition of 50 µM capsaicin. Each data point represents 1 female: N=58-60 females/meal (mean±SD). (E, G) Female engorgement on the indicated meal delivered via Glytube. Each data point denotes 1 trial with 15-20 females/trial: N=6-16 trials/meal. (F, H, J) Sampled weight measurements from engorged females offered the indicated meal or unfed controls not offered any meal from data in (E, G, I), respectively. N=3-25 weight measurements. Data labeled with different letters are significantly different from each other (mean±SEM; one-way ANOVA with Dunnett’s multiple comparisons with a single pooled variance, p < 0.05). (I) Female engorgement on the indicated concentration of ATP delivered in saline via Glytube. Each data point denotes 1 trial with 15-20 females/trial, N=4-14 trials/meal (mean±SEM). Ligands: saline = 110 mM NaCl and 20 mM NaHCO_3_; blood proteins = 15 mg/mL gamma-globulin, 8 mg/mL hemoglobin, 102 mg/mL albumin in 110 mM NaCl and 20 mM NaHCO_3_ (Duvall et al., 2019; Kogan, 1990); AMP-CPP (α,β-methyleneadenosine 5’-triphosphate lithium salt), AMP-PNP (β,γ-imidoadenosine 5’-triphosphate lithium salt hydrate), AMP-PCP (β,γ-Methyleneadenosine 5′-triphosphate disodium salt). (D,E,G) Data labeled with different letters are significantly different from each other (Kruskal-Wallis test with Dunn’s multiple comparison, p < 0.05).

**Figure S2:**
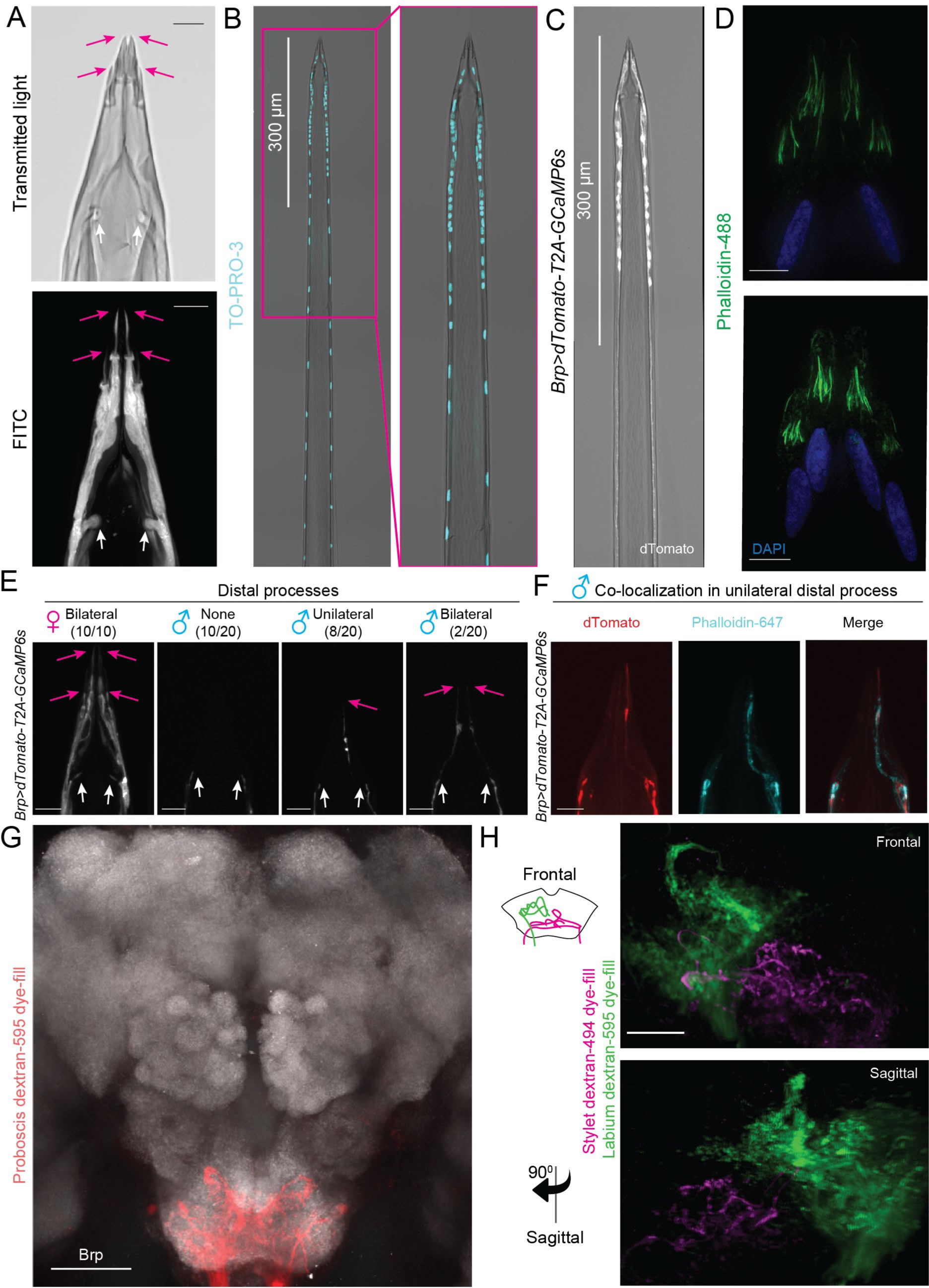
Sexual dimorphism in the stylet (Related to Figure 3) (A) Confocal images of transmitted light (top) and FITC counterstain (bottom) outline the female stylet chemosensory (pink arrows) and mechanosensory (white arrows) sensillar structure. (B,C) Tiled confocal image with transmitted light overlay of TO-PRO-3 nuclear staining (B, cyan) in a wild-type female stylet and dTomato expression (C, gray) in a *Brp*>*dTomato-T2A-GCaMP6s* female stylet. Right panel in (B) is a zoom-in of left panel and the same magnification is maintained in (C). (D) Super-resolution structured illumination image of phalloidin-488 actin stain (green) and DAPI nuclear stain (blue) in the female stylet tip from two wild-type individuals. Scale bar: 5 µm. (E) Confocal image of dTomato expression in the female (left) and male (remaining 3 panels) stylet tip of *Brp*>*dTomato-T2A-GCaMP6s* animals. From left to right: 10/10 females examined have extensive bilateral distal processes, 10/20 males examined have no distal processes, 8/20 males examined have sparse unilateral distal processes, and 2/20 males examined have sparse bilateral distal processes. (F) dTomato expression (left) and phalloidin-647 actin staining (middle) co-localize in the *Brp*>*dTomato-T2A-GCaMP6s* male stylet. Right panel is a merge of left and middle panel. (G) Proboscis neuron projection pattern (red) is restricted to the suboesophageal zone as revealed by dextran-595 dye fill. The proboscis consists of the stylet and labium, neuropil stained with anti-*Drosophila* Brp (gray). (H) Zoom-in on subesophageal zone after dual dye-fill with bilateral stylet dextran-494 (magenta) and unilateral labium dextran-595 (green). Bottom panel is a 90° optical rotation from the sagittal perspective. Scale bar: 10 µm (A,E,F), 25 µm (G,H) (See also Video 2).

**Figure S3:**
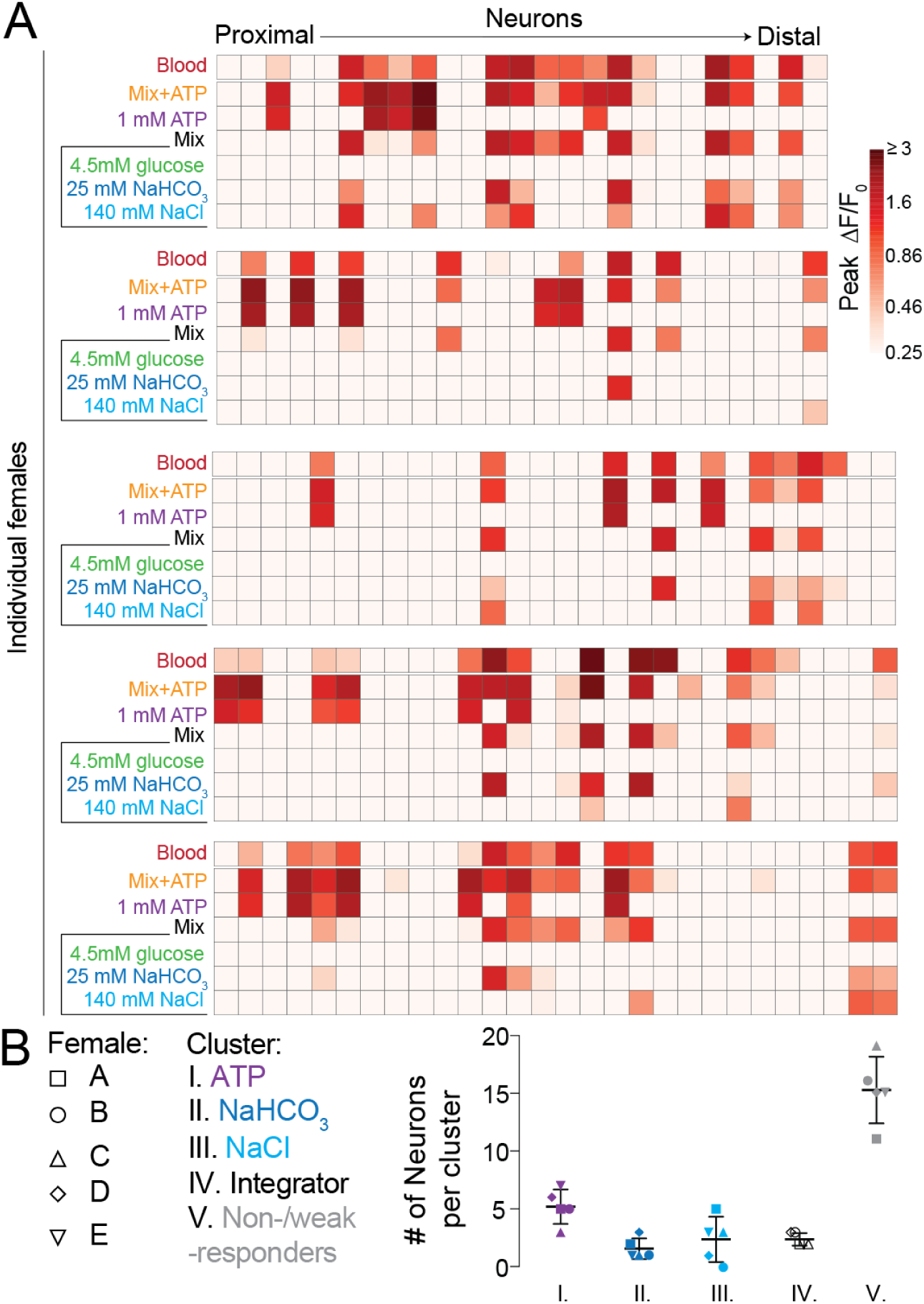
Blood component responses of individual females (related to Figure 5) (A) Heat maps of peak ΔF/F_0_ response to the indicated ligand for individual females prior to clustering in Figure 4G. Each square is the average of 3 ligand exposures. Each column represents one neuron and each row represents the response to indicated ligand for all neurons from 1 individual female. Neurons are ordered from proximal to distal. N=5 individual females. (B) Number of neurons per cluster in Figure 4G. All females have neurons in every cluster with the exception of Female D, which has neurons in all clusters except for the NaCl cluster.

**Figure S4:**
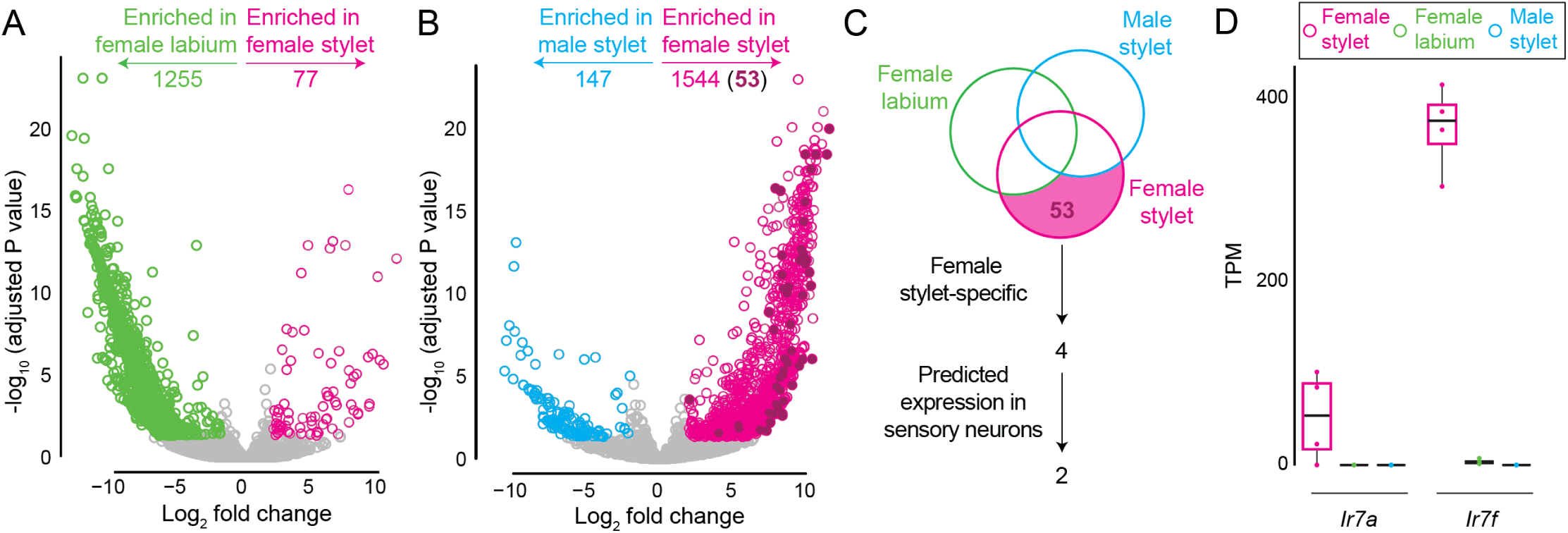
Female stylet-specific transcripts (Related to Figure 6): (A-D) RNA-seq dataset comparing the female stylet (pink), female labium (green), and male stylet (blue). N=4 replicates/tissue. (A,B) Volcano plot of transcripts enriched in the female stylet (pink) or female labium (green) in (A), and female stylet (pink) or male stylet (blue) in (B). 53 transcripts (fuchsia) were enriched in the female stylet compared to both female labium and male stylet. Transcripts were identified as significantly enriched in indicated tissue if Log_2_ fold change > 2 and adjusted p value < 0.05, as determined by DESeq2 differential expression analysis. (C) Venn diagram schematizing filters for identifying female stylet-specific transcripts. (D) Transcripts per million (TPM) expression data represented as box plots for putative female stylet-specific transcripts selected as driver lines. Median indicated by black line, bounds of box represent first and third quartile, whiskers are 1.5 times the inter-quartile range and outliers denoted by dot.

**Figure S5:**
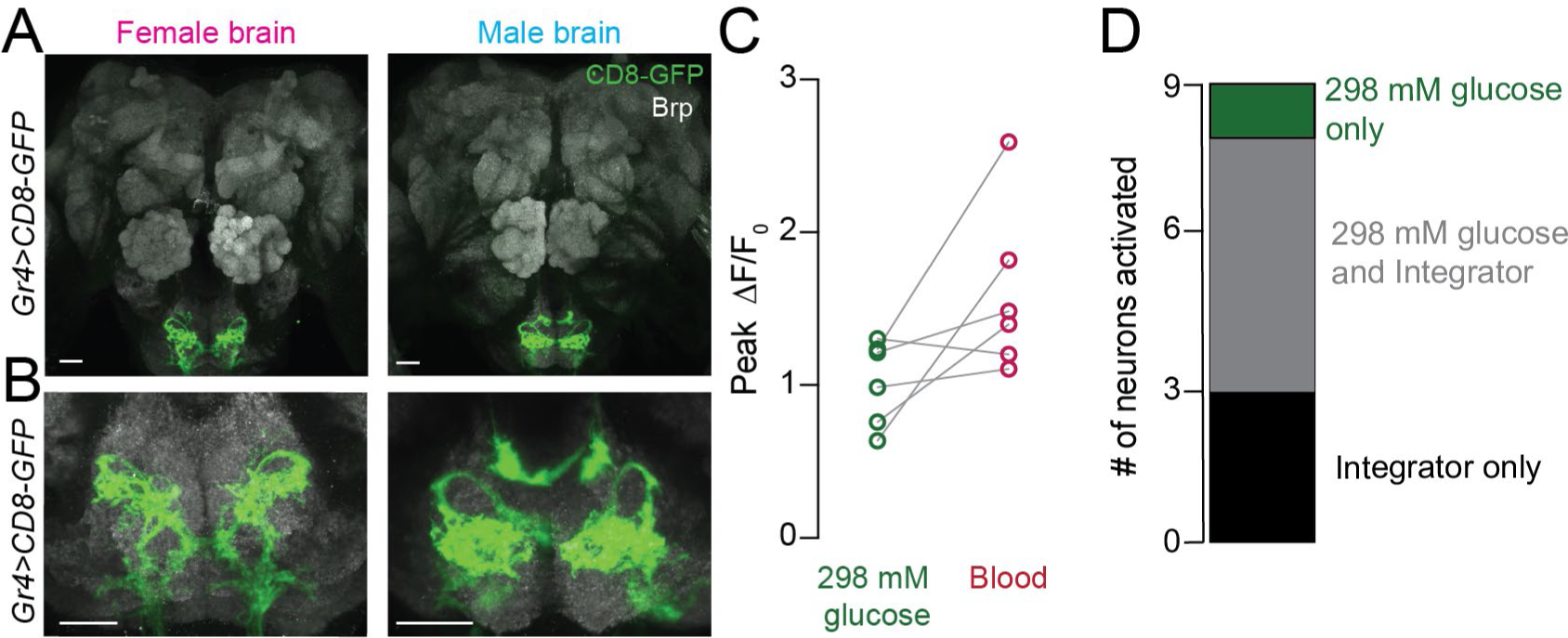
298 mM-sensitive neurons intersect with Integrator neurons (Related to Figure 7): (A,B) mCD8:GFP expression (green) of *Gr4*>*mCD8:GFP* in female (left) and male (right) brain (A) and subesophageal zone (B). Neuropil labeled with anti-*Drosophila* Brp (gray). Brain and subesophageal zone images were acquired from two different individuals. (C) For 298 mM-sensitive neurons (response to 298 mM glucose ≥ 0.25 peak ΔF/F_0_), peak ΔF/F_0_ to 298 mM glucose, compared to blood. Each data point denotes the response from 1 neuron and responses from the same neuron are connected by a line (N=6 neurons, mean±SD, p < 0.05, one-sample Wilcoxon signed rank test). (D) A dataset from N=5 females was filtered for all 298 mM-sensitive neurons and Integrator neurons in order to compare the intersection of 298 mM-sensitive neurons and Integrator neurons (N=9 neurons; 1/9 = 298 mM glucose only, 5/9 = 298 mM glucose and Integrator, 3/9 = Integrator only).

## Notes

https://github.com/VosshallLab/Jove_Vosshall_2020

